# Driving Osteocytogenesis from Mesenchymal Stem Cells in Osteon-like Biomimetic Nanofibrous Scaffolds

**DOI:** 10.1101/2022.06.28.497866

**Authors:** Farhad Soheilmoghaddam, Hadi Hezaveh, Madeleine Rumble, Justin J. Cooper-White

**Affiliations:** Tissue Engineering and Microfluidics Laboratory, The Australian Institute for Bioengineering and Nanotechnology (AIBN), University of Queensland, St. Lucia, QLD, 4072, Australia; School of Chemical Engineering, University Of Queensland, St Lucia, QLD, 4072, Australia

**Keywords:** Bilayered scaffolds, BMP-6 peptide, Smad signalling pathway, osteocyte-like cells

## Abstract

The repair of critical-sized bone defects, resulting from tumor resection, skeletal trauma or infection, remains a significant clinical problem. A potential solution is a tissue-engineered approach that utilises the combination of human mesenchymal stem cells (hMSCs) with synthetic biomaterial scaffolds, mimicking many of the biochemical and biophysical cues present within the native bone. Unfortunately, osteocyte cells, the orchestrators of bone maturation and homeostasis, are rarely produced within such MSC-seeded scaffolds, limiting the formation of true mature cortical bone from these synthetic implants. In this contribution, a bone morphogenic protein-6 (BMP6)-presenting osteon-like scaffolds based on electrospun poly(lactic-co-glycolic acid) (PLGA) fibrous scaffolds and poly(ethylene glycol) (PEG) based-hydrogels is reported. BMP6 peptide is shown to drive higher levels of SMAD signalling than the full-length protein counterpart. Osteon-mimetic scaffolds promoted the formation of osteocyte-like cells displaying multi-dendritic morphology and osteocyte-specific marker, E11/gp38 (E11), along with significant production of dentin matrix protein 1 (DMP1), confirming maturation of the ososteocyte-like cells. These results demonstrate that osteon-like scaffolds presenting chemo-topographical cues can drive the formation of mature osteocyte-like cells from hMSCs, *without* the need for osteogenic factor media supplements, providing a novel ex vivo production platform for osteocyte-like cells from human MSCs in cortical bone mimics.

## 1. Introduction

Bone repair is a complex cascade of biological events that depend on the coordination of mesenchymal stem cells (MSC), adhesion, proliferation and osteogenic differentiation. ^[1]^. During this process, complex interplays between multiple signalling pathways, including, critically, bone morphogenic proteins (BMPs), drive the differentiation of recruited MSCs cells that reside within bone marrow into osteoblasts (bone-forming cells) and thereafter osteocytes (mature bone-regulatory cells)^[2]^ that participate in the creation and maturation of new cortical bone, providing bone with an enviable regenerative capacity compared to other tissues – *if the defects are small*.

In the case of critical-sized defects (>1–3 cm in size), this highly orchestrated and ‘encoded’ process, in which MSCs are first recruited to the injury site before undergoing stagewise differentiation into mature bone-regulatory cells, is insufficient for ‘bridging the gap’, requiring the assistance of a matrix (e.g. bone graft) or scaffold to capitulate with the endogenous (or exogenous) MSCs to ‘engineer’ new bone in the defect ^[3]^. Bone grafts, including autogenous or allogenic materials, are currently the clinically preferred choice for bone repair, but the shortage of autograft tissue, risks of disease transfer and histoincompatibilities ^[4]^, and mechanical-mismatch driven adjacent tissue degeneration (in the case of metallic cages) ^[5]^ are just some of their limitations. These limitations have prompted researchers in the bone tissue engineering field to explore new approaches to overcome these roadblocks to clinical translation, including most recently, the incorporation of biochemical and biophysical cues provided in native cortical bone during repair into engineered scaffolds, especially those present in the osteon, the ‘functional unit’ of cortical bone ^[6]^.

Biochemical cues in such scaffolds are provided by a multitude of growth factors, differentiation factors or drugs. Of these, Bone Morphogenic Proteins (BMPs), multifunctional growth factors within the transforming growth factor β (TGF-β) superfamily, are most frequently used due to their unique osteoinductive potential and crucial roles in skeletal development and maintenance ^[7]^. The canonical BMP signaling cascade initiates when BMPs associate with type I and type II BMP receptors to form a multimeric receptor ligand complex, followed by phosphorylation of BMP-specific Smads, including Smad 1, 5 and 8, to regulate the expression of the targeted genes, including Runx 2 and ALP ^[7c, 8]^. Of the known BMP variants, BMP2, BMP6 and BMP9 have been shown to be the most potent inducers of osteoblast differentiation from MSCs ^[7e, 9]^. Of these, although BMP2 has received FDA approval for clinical use ^[10]^, BMP6 has been shown to be more potent (both *in vitro* and *in vivo*) ^[11]^. Regardless, despite the remarkable potential of BMPs for bone regeneration and repair, broader clinical application is often hindered by their random folding, short half-life, high prices and immunogenicity ^[4b, 11a, 12]^. A viable alternative is a short peptide of the active domains of the protein capable of replicating the signaling cascades induced by the full length protein. Studies have shown that the biological activity of BMP-derived peptides is highly influenced by their carriers and that carriers designed to sustain the bioactivity and release of BMP peptides can lead to reduced effective dose sizes and reduced occurrence of side effects compared to full length proteins ^[13]^.

Biophysical cues have been incorporated into such scaffolds using a multitude of architectural and morphological mimicry of the native bone microenvironment. In particular, many studies have proposed the use of electrospun nanofibres for synthetic bone grafts due to their resemblance (in lengthscales and architecture) to native bone ECM structures and their potential to be equipped with BMP proteins or peptides ^[13b, 14]^. None of these works, however, have successfully created scaffolds that replicate both the micro and nanostructural organisation of an osteon ^[6c, 15]^. Moreover, these works are yet to confirm effective *in vitro* differentiation of MSCs in such scaffolds to *bonafide* osteocytes that display phenotypic morphological features and marker expression (including transmembrane glycoprotein E11, expressed in newly embedded osteocytes), and dentin matrix acidic phosphoprotein 1 (DMP1, required for osteocyte maturation) ^[16]^, whilst *also* better matching topographical and mechanical property requirements of cortical bone.

Recently, we ^[17]^ utilised electrospinning approaches as a simple and scalable method of fabricating aligned nanofibers of PLGA encapsulating rod-shaped nano-sized hydroxyapatite (HA) (PLGA-HA). By encoding axial orientation and spacing of the HA within these aligned nanocomposite fibres, significant improvements in a mineral deposition under simulated body fluid (SBF) conditions and in mechanical properties (in the axial direction) were achieved. For the first time, we showed that when osteon-inspired chemical and topographical cues are provided to MSCs in our electrospun mats, pro-osteogenic factors (e.g. dexamethasone) were not required in the media to drive osteoblastogenesis. However, these cells did not encase themselves in osteoid, and hence osteocytogenesis was not observed. Clearly, the chemo-topographical cues from these aligned HA spatially encoded nanofibers alone were not enough.

In this work, to better mimic the microenvironmental architectural and cytokine cues existing in the osteon, hBMSCs were seeded onto a bilayered construct composed of aligned nanofibrous PLGA-HA-based scaffolds and subsequently *overlayed* with a hydrogel activated with a synthesised short peptide mimic of the receptor-binding domain of BMP-6 protein. An outline of the conducted experiments is schematically represented in **Figure 1**. Panel A: First, the bilayered scaffold was constructed after optimisation of each of the fabrication processes. Electrospinning was utilised to fabricate aligned PLGA bioactive electrospun fibres reinforced with highly oriented nHA that mimic the native bone in terms of ECM niche and matrix structure. This outcome was obtained by tuning the polymer concentration, nHA morphology and the electrospinning parameters, with the optimum detailed in ‘Methods’. hBMSCs were cultured on mono-layered scaffolds in media with different compositions including growth medium (GM), complete osteogenic medium (OS) and OS medium in the absence of DEX (OS-DEX). Panel B: Second, to enable sustained bioactivity of BMP6-derived peptides, the cell-seeded PLGA-HA scaffold was combined with a layer of BMP6 peptide-non-conjugated and -conjugated PEG hydrogel (using norbornene-thiol chemistry). An optimal dosage of BMP6 peptide based on P-Smad 1/5/8 expression results was used for the fabrication of the bilayered scaffold. As we have shown previously ^[18]^, the biological activity of many short chain molecules (peptides, GAGs, etc.) are best sustained and more potent when covalently bound to macromolecular scaffolds (such as biopolymers). Thus, we employed thiol-end click chemistry to conjugate our BMP6 peptide to polyethylene glycol (PEG)-star polymer (BMP6p-PEG). The BMP6p-PEG hydrogel precursor was introduced as a thin coating on top of hBMSCs pre-seeded on the PLGA-HA electrospun fibrous scaffold. By combining the confirmed osteoconductive and inductive cues of the aligned nanofibrous HA-loaded scaffold with the inductive cues provided by localised and sustained BMP6 growth factor signalling, we aimed to assess the ability of these hierarchical scaffolds to promote osteocyte formation and mature bone production from hBMSCs. Our results confirm that the provision of biochemical and architectural cues to hBMSCs in spatially controlled niche-mimicking environments present during bone repair drives the formation of human osteocyte-like cells and supports neo-bone tissue formation under *in vitro* culture conditions, even in standard hBMSC maintenance media conditions.

**Figure 1.**
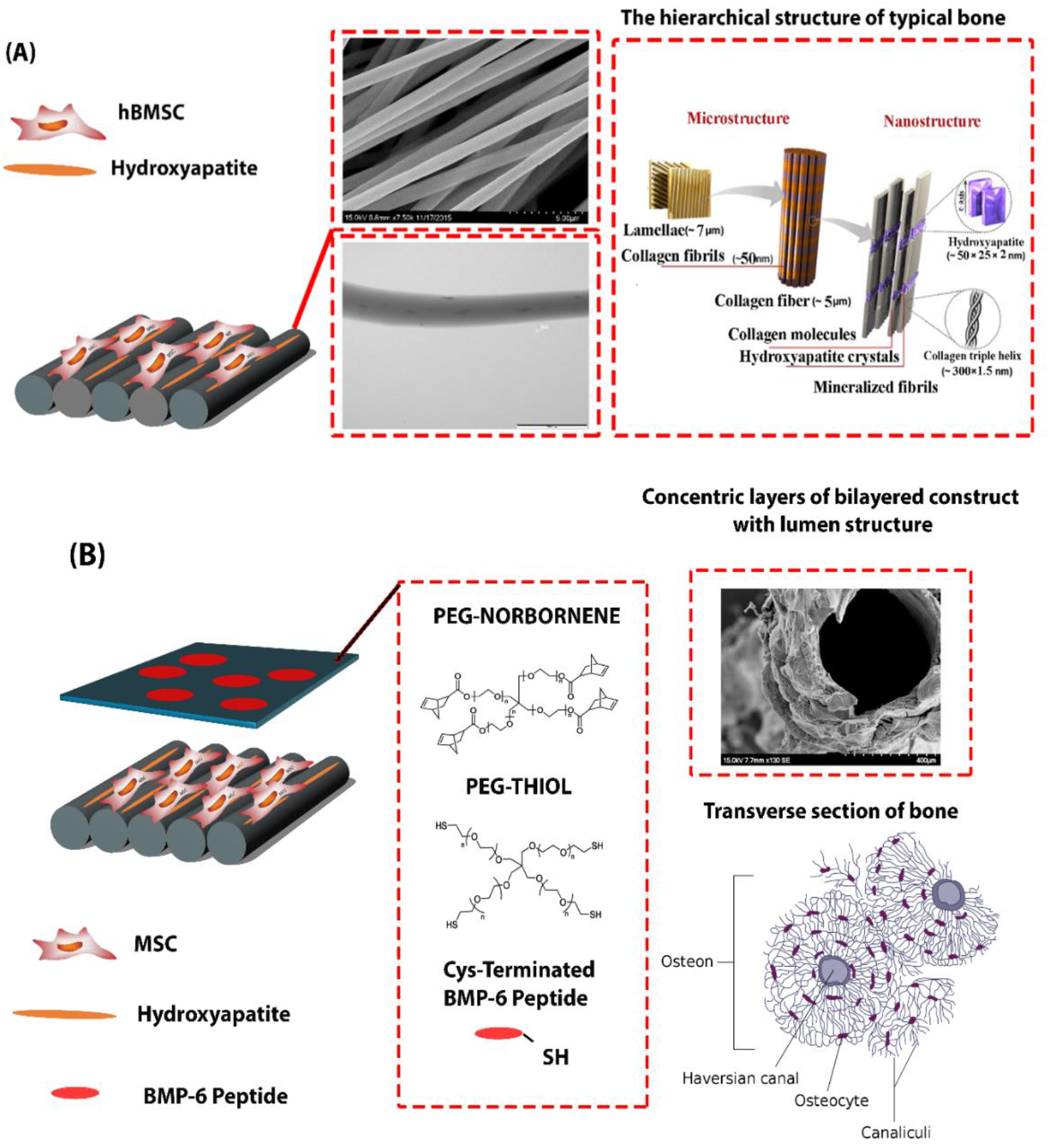
Panel A: Schematic illustration of aligned PLGA electrospun fibres reinforced with rod-shape nHA mimicking the ECM structure of native bone. Panel B: Schematic representation of bilayered scaffold comprised of electrospun nanofibres and BMP6 peptide-conjugated PEG hydrogel. The hierarchical structure of typical bone was retrieved from ^[45]^ and the transverse section of bone from: https://commons.wikimedia.org/wiki/

**Figure 2.**
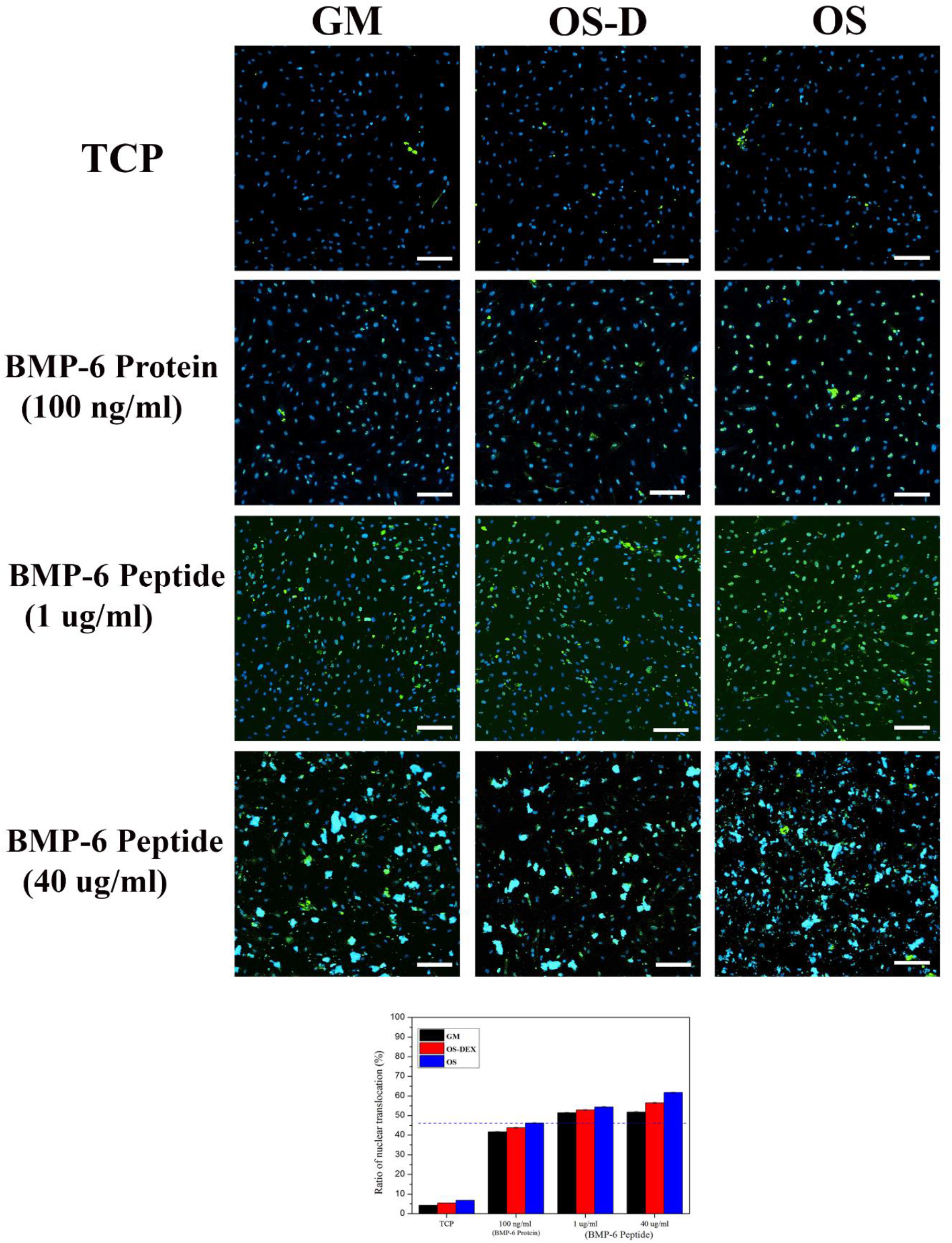
Immunohistochemistry expression of P-Smad 1/5/8 after 3 hourrs treatment of hBMSCs with different dosages of BMP6 peptide in GM, OS-DEX an OS. P-Smad 1/5/8 are labelled in green and the cell nuclei are labelled in blue. Scale bars, 200 μm; *P<0.05.

## 2. Results and Discussion

### 2.1. Validation of BMP6-derived peptide activity and unction

BMP6 proteins, like other members of the TGF-β superfamily, bind to both type I and type II receptors, but unlike the TGF-β superfamily, BMP6 binding affinity is upregulated in the presence of both receptor types ^[19]^. The phosphorylated receptor induces the Smad 1/5/8 phosphorylation, leading to the formation of heteromeric complexes with Smad4, which translocate into the nucleus ^[20]^. To confirm that the synthesised BMP6 peptide was active and functional via these expected pathways, the nuclear localisation of phosphorylated Smad1/5/8 (P-Smad 1/5/8) in hBMSCs cultured with different dosages of BMP6 peptide (added to media) was evaluated after 24 hours (**Figure S1**). Using 100 ng/ml of BMP6 protein first as the positive control, P-Smad 1/5/8 nucleus localisation was abundant when the cells were exposed to 1 μg/ml of BMP6 peptide or higher dosage, while significant cytoplasmic localisation of P-Smad 1/5/8 was also observed when the cells were exposed to 40 ug/ml of BMP6 peptide. Quantification of the level of nuclear localisation of P-smad 1/5/8 confirmed active BMP6 signalling pathway at dosage of 1μg/ml or greater. At these doses, signalling was equal to or higher that the positive control.

hBMSCs respond very rapidly to BMP signalling ^[21]^ and thus, the P-Smad 1/5/8 expression of hBMSCs cultured with BMP6 protein and peptides was examined shortly after incubation (3 hours) (**Figure 3**). Higher levels of P-Smad 1/5/8 were observed 3 hours after incubation compared to similar samples that were incubated for 24 hourrs. This suggests the rapid phosphorylation of Smad 1/5/8 is induced by the BMP6 peptide. The level of P-Smad 1/5/8 expression for hBMSCs incubated for 3 hours with 40 ug/ml BMP6 peptide was less than at the longer culture time (24 hours), whilst the cytoplasmic expression of P-Smad 1/5/8 also decreased in the shorter incubation time. The substantial cytoplasmic localisation of P-Smad 1/5/8 for cells exposed to high dosages of BMP6 peptide after 24 hours is thus likely due to rapid saturation of the BMP receptors on hBMSCs at such high BMP6 peptide dosage, leading to dephosphorylation of P-Smad 1/5/8 and cytoplasmic localisation. The results for cytoplasmic localisation of P-Smad 1/5/8 at high doses of BMP6 peptide are in agreement with those of Chen and co-workers ^[22]^, who probed the BMP-dependent mouse MSCs differentiation activity and transcriptional responses into the BMP2 protein.

**Figure 3.**
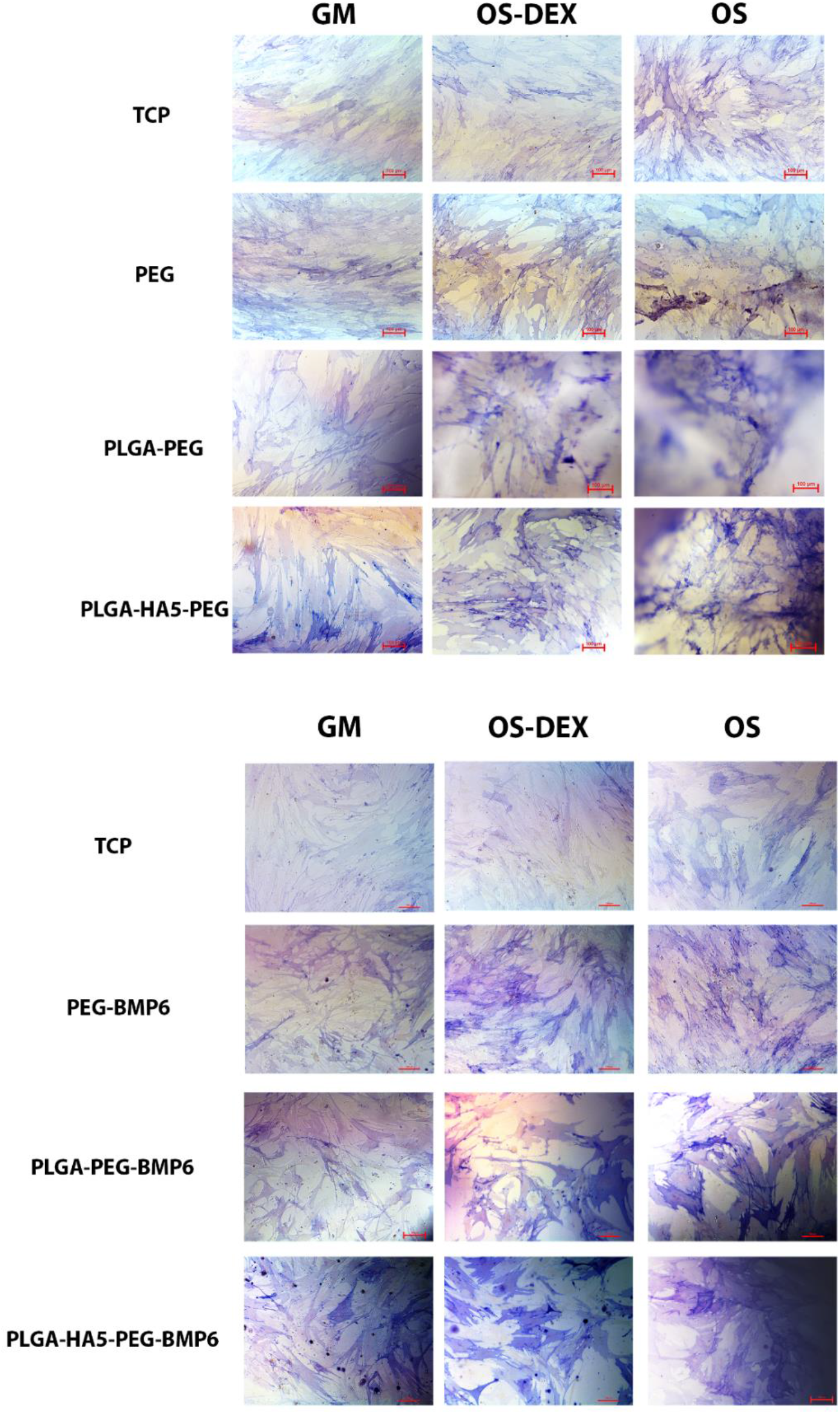
ALP staining of hBMSCs cultured on TCP, PLGA and PLGA-HA5 in the presence of non-conjugated PEG hydrogel or BMP6 conjugated hydrogel in different medium conditions for 7 days.

It is clear from the P-Smad 1/5/8 expression in different medium conditions that DEX can increase Smad 1/5/8 phosphorylation ^[23]^. However, regardless of medium conditions, a high level of P-Smad 1/5/8 expression was obtained when cells were exposed to a high dosage of BMP6 peptide (1 μg/ml), indicating the high affinity of the synthesised peptide for the receptor and its ability to activate the Smad signaling pathway.

#### 3.4.2. Effect of different dosage of synthesised BMP6 peptide on osteogenic differentiation of hBMSCs in different media conditions

Next, we utilised thiol-end click chemistry to conjugate the BMP6 peptide (terminated with a cysteine) to the backbone of the PEG hydrogel (BMP6p-conjugated PEG) that will cover the hBMSC layer, once seeded onto the electrospun scaffold (Panel B). Before investigating the influence of the BMP6 peptide on the osteogenic differentiation of hBMSCs, the cell viability in the presence of PEG hydrogel was first assessed (**Figure S2**). The live/dead assay results of cultured hBMSCs on TCP with a layer of PEG hydrogel on top showed that the cell morphology remained unchanged and hBMSCs were viable after 24 hours of incubation in different medium conditions.

The ability of osteoinductive molecules to induce bone regeneration is highly dependent on the biophysical and architectural properties of their carrier. To investigate this, we characterised the mechanical and morphological characteristics of the PEG-Nor/PEG-TH hydrogels. The rheological and morphological properties of PEG-Nor/PEG-TH hydrogels with 4% (w/v) and 10% (w/v) of polymer content was assessed. The PEG-Nor/PEG-TH hydrogel with 4% (w/v) polymer content showed the storage (elastic) modulus (*G*′) of 4kPa and loss modulus (viscous) (G″) of 21Pa with pore sizes ranging from 120 nm to 300 nm. When the polymer concentration increased to 10% (w/v), *G*′ increased by two times (8 kPa), as expected while the (G″) was increased to 47Pa. The Cryo-SEM images of this hydrogel show interconnected pores, with sizes ranging from 26 nm to 120 nm. The BMP6 conjugated PEG-Nor/PEG-TH hydrogel with 10% (w/v) polymer concentration showed slightly higher *G*′ (11 kPa) when compared to the non-conjugated PEG-Nor/PEG-hydrogels (**Figure S3**).

To assess whether the BMP6 peptide, when conjugated to the PEG hydrogels, can still induce Smad signaling, the peptide was added to the medium during crosslinking of the gel. P-Smad 1/5/8 expression of the hBMSCs cultured with BMP6-PEG with 1 μg/ml of BMP6 was investigated after 24 hours. A high level of P-Smad 1/5/8 nuclei localisation was observed for hBMSCs cultured in different medium conditions. Quantitative analysis of P-Smad 1/5/8 expressed in nuclei showed slightly higher expression for hBMSCs cultured with BMP6 conjugated hydrogels compared to when hBMSCs were incubated with a similar concentration of free BMP6 peptide (i.e. simply added to the medium) (**Figure S4**). This suggests that the BMP6 conjugated PEG hydrogel was able to sustain the bioactivity of the BMP6 peptide with even higher affinity compared to the full length BMP6 protein, at a reduced peptide dosage. As we have shown previously, the biological activity of many short-chain molecules (peptides, GAGs, etc.) ^[18]^ are best sustained and more potent when covalently bound to macromolecular scaffolds (such as biopolymers). For the remaining experiments, we selected the following composition: PEG hydrogel composition with 10% (w/v) and the 1 μg/ml BMP6 peptide.

To support the immunofluorescence staining results, the expression of Smad 1/5/8 genes were measured using qPCR (**Figure S5**). Compared to the hBMSCs treated with BMP6 protein or peptide in medium, the BMP6 conjugated PEG sample presented slightly higher levels of Smad 1, 5 and 8. Highest expression of Smad 1, 5 and 8 genes in all medium conditions were observed when hBMSCs were exposed to 40 μg/ml of BMP6 peptide, indicating significant osteoinductive potential of the synthesised BMP6 peptide.

ALP expression, an early osteogenic marker, was next assessed for the hBMSCs exposed to different dosages of free BMP6 peptide incorporated into the medium compared with the PEG-BMP6 sample. Similar levels of ALP staining were observed for the cells cultured with 1 μg/ml of free BMP6 peptide added to medium compared to 100 ng/ml of the full length BMP6 protein. Interestingly, the BMP6-conjugated PEG samples exhibited the highest ALP staining compared to positive controls in similar medium conditions (**Figure S6**). The conjugation of BMP6 peptide to PEG-Nor/PEG-TH hydrogels is a promising strategy for sustaining the bioactivity of BMP6 peptides and supporting osteogenic differentiation.

ECM mineralisation is another critical indicator for hBMSCs osteogenic differentiation ^[24]^. To confirm the functionality of BMP6 peptide as an osteoinductive factor, the calcium deposition of hBMSCs exposed to different dosages of free BMP6 peptide in media and PEG-BMP6 was investigated (**Figure S7**). The Alizarin red staining showed that the BMP6 peptide alone significantly induced calcium deposition, in comparison to the BMP6 protein. It should be noted that a high level of calcium deposition was observed for BMP6-conjugated PEG hydrogel samples, even in medium *lacking* DEX or any osteogenic supplements, indicating the BMP6-conjugated PEG maintains BMP6 peptide bioactivity. Quantitative analysis of calcium deposition showed slightly higher calcium deposition for hBMSCs exposed to 1 μg/ml of BMP6 peptide compared to 100 ng/ml of BMP6 protein. Calcium deposition from cells covered with the BMP6-conjugated PEG hydrogel was significantly higher than those treated with 100 ng/ml of BMP6 protein or BMP6 peptide of similar dosage added into the medium. Moreover, the BMP6p-conjugated PEG hydrogels deposited higher levels of calcium in OS-DEX medium, as compared to TCP in complete OS medium.

Collagen 1 (Col 1) is a critical protein in the formation of new bone. We thus next investigated production of Col 1. In this experiment, hBMSCs cultured with 100 ng/ml of BMP6 protein in different medium condition was used as the positive control. The BMP6 peptide, when added to the medium, induced Col1 secretion after 7 days (**Figure S8)**. Compared to the BMP6 protein, the BMP6p-conjugated PEG hydrogels expressed higher levels of Col 1, further confirming these conjugated PEG hydrogels can sustain the bioactivity of BMP6 peptides to drive Col 1 production.

To validate the effect of BMP6 peptide on osteogenic maturation of hBMSCs, the production of late osteogenic protein markers osteopontin (OP) (**Figure S9**) and osteocalcin (OC) (**Figure S10**), was assessed. The results from immunofluorescence staining of OP and OC suggest that BMP6 peptide alone can promote maturation by driving high levels of OP and OC protein production. Additionally, when the hBMSCs were cultured with BMP6p-conjugated PEG hydrogel, staining for both OP and OC was high even in OS-DEX medium or GM. Notably, OP and OC production by hBMSCs cultured with BMP6p-conjugated PEG hydrogels in OS-DEX was higher than cells that were cultured with BMP6 protein in complete OS media. This suggests that the BMP6 conjugated PEG hydrogels can maintain the BMP6 peptide bioactivity even after long culture times, supporting osteoblast maturation (up to 21 days).

### 2.2. Osteogenic differentiation of hBMSCs seeded bilayered electrospun fibers in different media conditions

Osteogenic differentiation and maturation of hBMSCs on a bilayered scaffold, that combined the PLGA-HA electrospun scaffold with the BMP6p-conjugated PEG hydrogel, in the absence of DEX or any osteogenic supplements, was next assessed. Based on previous results surveying different dosages of BMP6 peptides, 1 μg/ml of BMP6 peptide was considered the optimal dosage for conjugation into the PEG hydrogels for further experiments.

ALP activity is critical in the early stages of hBMSC osteogenic differentiation. A high level of ALP staining was observed for the samples cultured with BMP6p-conjugated PEG hydrogels in all medium conditions (**Figure 3**). Interestingly, ALP expression for the PLGA-HA5-PEG-BMP6 sample in GM was significantly higher than those cultured on TCP in complete OS medium. It is clear that there were significant differences between the ALP expression in the samples cultured with BMP6p-conjugated PEG hydrogels compared to controls. However, control samples showed slightly higher positive staining when compared to TCP samples, suggesting that the ability of 3D PEG hydrogels to constrain or capture the secreted ECM proteins provides advantage in terms of supporting ALP production. We previously reported that these PEG-Nor/PEG-TH hydrogels are capable of capturing different ECM proteins such as fibronectin, laminin and Col1 ^[25]^.

Calcium deposition in the PLGA-PEG and PLGA-HA5-PEG scaffolds increased by almost 1 fold and 1.5 fold, respectively, when compared to TCP in complete OS medium (**Figure 4**), attributed to hBMSCs responding to topographical cues of the electrospun scaffolds with native bone mimicking features. A substantial amount of calcium was deposited on the samples that were cultured with BMP6p-conjugated PEG hydrogels in all medium conditions, due to the PEG hydrogel’s ability to sustain the bioactivity of BMP6 peptide. It is worth noting that the PLGA-HA5-PEG-BMP6 samples in OS-DEX expressed significantly higher calcium content compared to TCP in complete OS medium. This suggested the bilayered constructs are able to induce ECM mineralisation (Col 1, OP and OC), even in the absence of DEX or any osteogenic supplementation.

**Figure 4.**
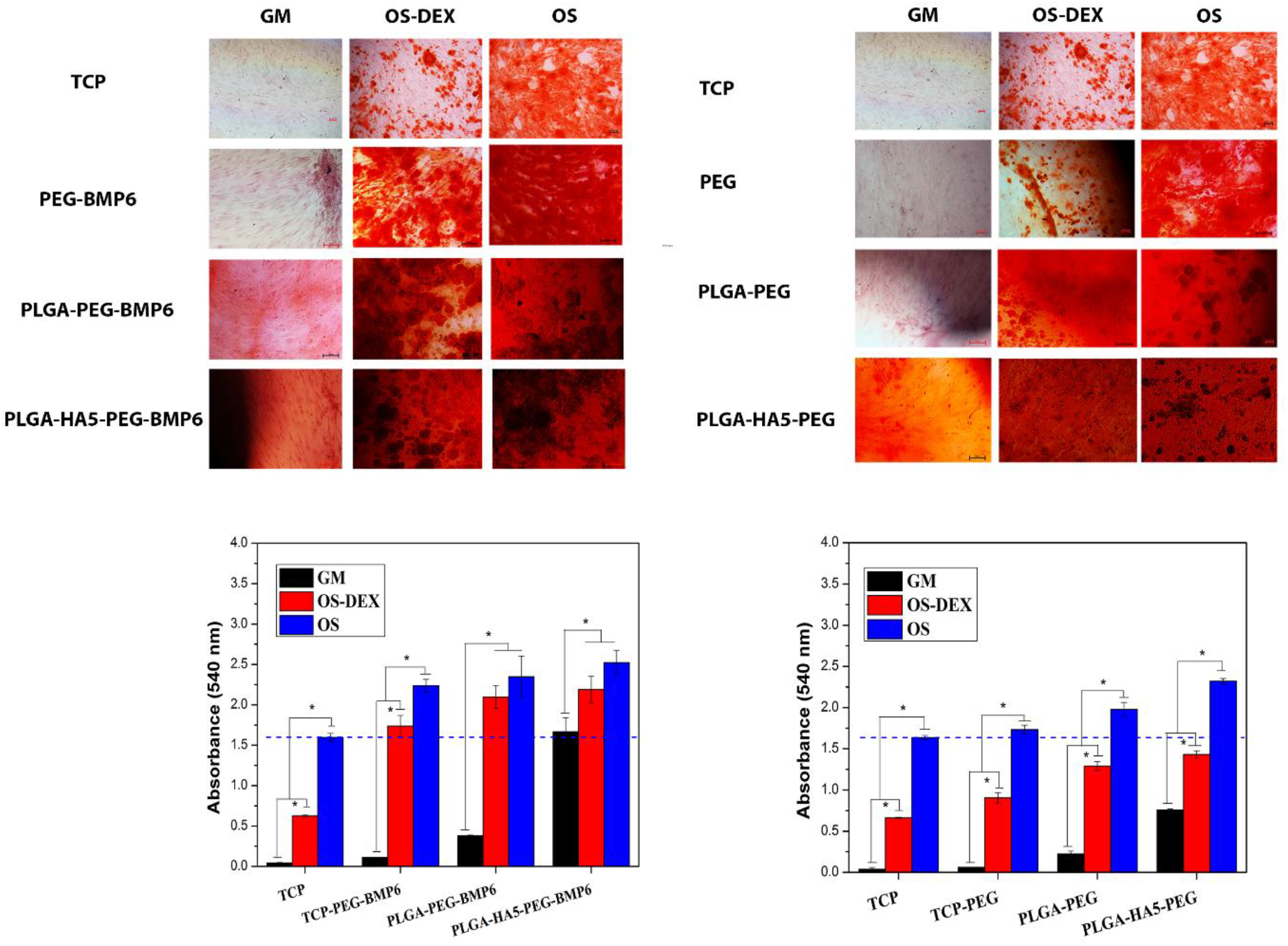
Alizarin red staining and quantified Alizarin red intensity after 21 days on the bilayered scaffolds incorporated with and without BMP6 peptide. Scale bar = 100 μm; *p < 0.05.

Following the calcium deposition analysis of the bilayered scaffolds by Alizarin red staining, HA formation was further assessed using on Osteo-Image mineralisation assay (**Figure S11**). Similar to calcium deposition results, there was no notable HA on TCP except in complete OS medium. The inclusion of the PEG hydrogel slightly increased HA formation as compared to TCP, which may be attributed to the ECM protein capturing potential of these PEG hydrogels. The whole surface of the PLGA-PEG-BMP6 scaffold was covered with small HA particles in OS medium, whilst in the case of the PLGA-HA5-PEG-BMP6 scaffold, it induced substantial HA formation, even in OS-DEX medium. Large HA aggregates were observed in the PLGA-HA5-PEG-BMP6 samples, suggesting the formation of bone-like nodules, even in GM. These results have further emphasised the potential of these bilayered scaffolds to promote bone mineralisation in the absence of osteogenic supplements to culture media.

To further assess the morphological properties and chemical composition of the minerals that were produced during differentiation of hBMSCs on these bilayered scaffolds, we utilised SEM and EDX analysis (**Figure S12**). The SEM images of mono-layered PLGA-HA5 samples seeded with hBMSCs showed the surface of the nanofibrous scaffolds were completely covered with rough multi-layers of cells, ECM and mineral. The hBMSCs on PLGA-HA5 scaffolds displayed a spindle-like morphology, sending out their projections to the fibre layers, suggesting strong cell–matrix interation (**Figure S12 a and b**). The hBMSCs on the PLGA-HA5-PEG scaffolds displayed an osteoblastic morphology (**Figure S12 c and d**) and secreted apatite-like crystals (**Figure S12 h**). The EDX analysis of deposited minerals confirmed the apatite formation was composed of calcium and phosphate ions, with the ratio similar to native bone apatite. After BMP6 conjugation into the PEG hydrogel, cell morphological characteristics substantially changed with the formation of dendritic processes that were greater in length than the cell body (**Figure S12 g and h**), suggesting osteocyte formation ^[26]^. Interestingly, the cell body size of the osteocyte-like cells varied between 5 to 15 microns, which is very close to native human bone osteocytes ^[27]^.

We next investigated the expression of Col1, the most abundant organic component of bone matrix. The bilayered scaffolds in the absence of BMP6 peptide produced high levels of Col1 when cultured in compelete OS medium for 7 days. (**Figure 5**a). Prior to the incorporation of BMP6 peptide into the samples, very low levels of Col1 were observed in OS-DEX or GM. The PLGA-HA5-PEG scaffolds were the only samples that induced Col1 production in OS-DEX medium or GM after only 7 days (**Figure 5**a), attributed to the topographical cues of the PLGA-HA5-PEG scaffolds, promoting osteoblastic differentiation of hBMSCs ^[17]^.

**Figure 5.**
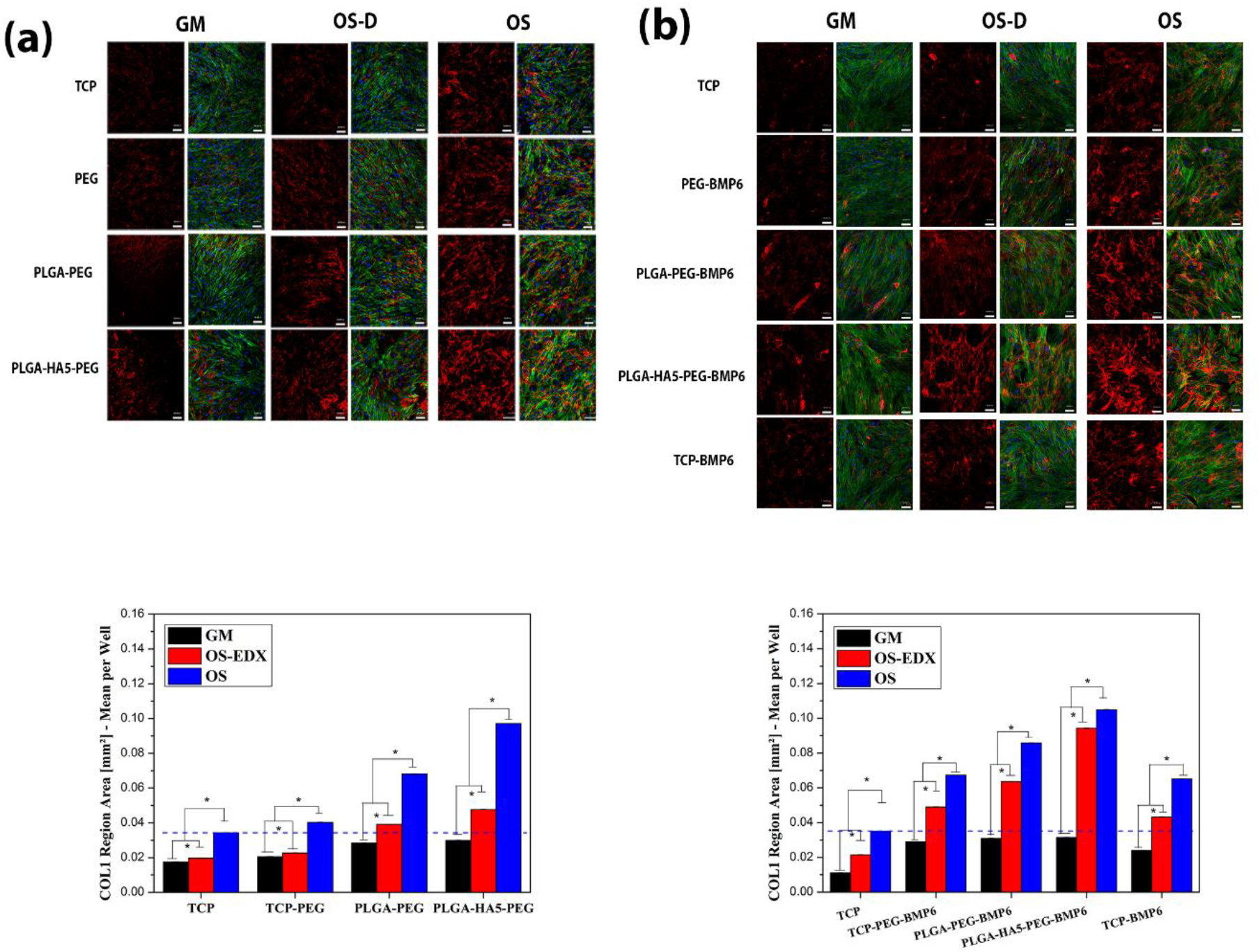
Immunofluorescence staining of Col1 for samples cultured with PEG hydrogels (a) before and (b) after BMP6 conjugation. Col1 expression was quantified based on immunofluorescence staining pictures. Scale bars, 200 μm; *P<0.05.

Thereafter, the impact of BMP6p-conjugation to the PEG hydrogel on Col1 production was assessed (**Figure 5**b). The BMP6p-conjugated PEG hydrogel induced substantial Col1 production. Quantification of these results showed significantly higher levels of Col1 on both PLGA-PEG-BMP6 and PLGA-HA5-PEG-BMP6 scaffolds in OS-DEX and GM medium, when compared to TCP in complete OS medium. Interestingly, although the hBMSCs cultured with free BMP6 peptide in the medium were exposed to three times higher amounts of BMP6 peptide (1μg/ml of BMP6 peptide was added to the medium, while the cells were fed every 2– 3 days), compared to the bilayered scaffolds with conjugated BMP6 peptide, the bilayered scaffolds displayed higher levels of Col1 protein. This further confirmed the ability of these novel bilayered scaffolds to support the bioactivity of BMP6 peptide, leading to substantial ECM production and osteogenic induction at early stages of differentiation.

Maturation of osteoblast cells was further assessed after 21 days, using immunofluorescence staining of OP (**Figure 6**) and OC proteins (**Figure 7**). Both PLGA-PEG and PLGA-HA5-PEG samples produced significantly higher levels of OP as compared to TCP in complete OS medium (**Figure 6**a). However, after the incorporation of BMP6p-conjugated PEG hydrogel, the bilayered scaffolds produced substantially higher OP levels compared to the TCP sample in all medium conditions (**Figure 6**b). Interestingly, the OP produced on bilayered scaffolds in GM was significantly higher than the TCP in complete OS medium. The bilayered scaffolds, especially PLGA-HA5-PEG-BMP6, induced substantial levels of OC protein (**Figure 7**a). There was a significant difference in OC expression of PLGA-HA5-PEG-BMP6 in GM, lacking any osteogenic supplements compared to TCP in OS medium. Notably, regardless of medium conditions, the BMP6 conjugated PEG hydrogels expressed significantly higher OP and OC proteins compared to the samples that were cultured with free BMP6 peptide in the same medium. It should be noted that the hBMSCs on bilayered scaffolds were exposed to 10 times less BMP6 peptide compared to the cells treated with free BMP6 peptide in medium after 21 days; every 2 days the cells treated with free BMP6 peptide were fed with medium containing 1 μg/ml of BMP6 peptide.

**Figure 6.**
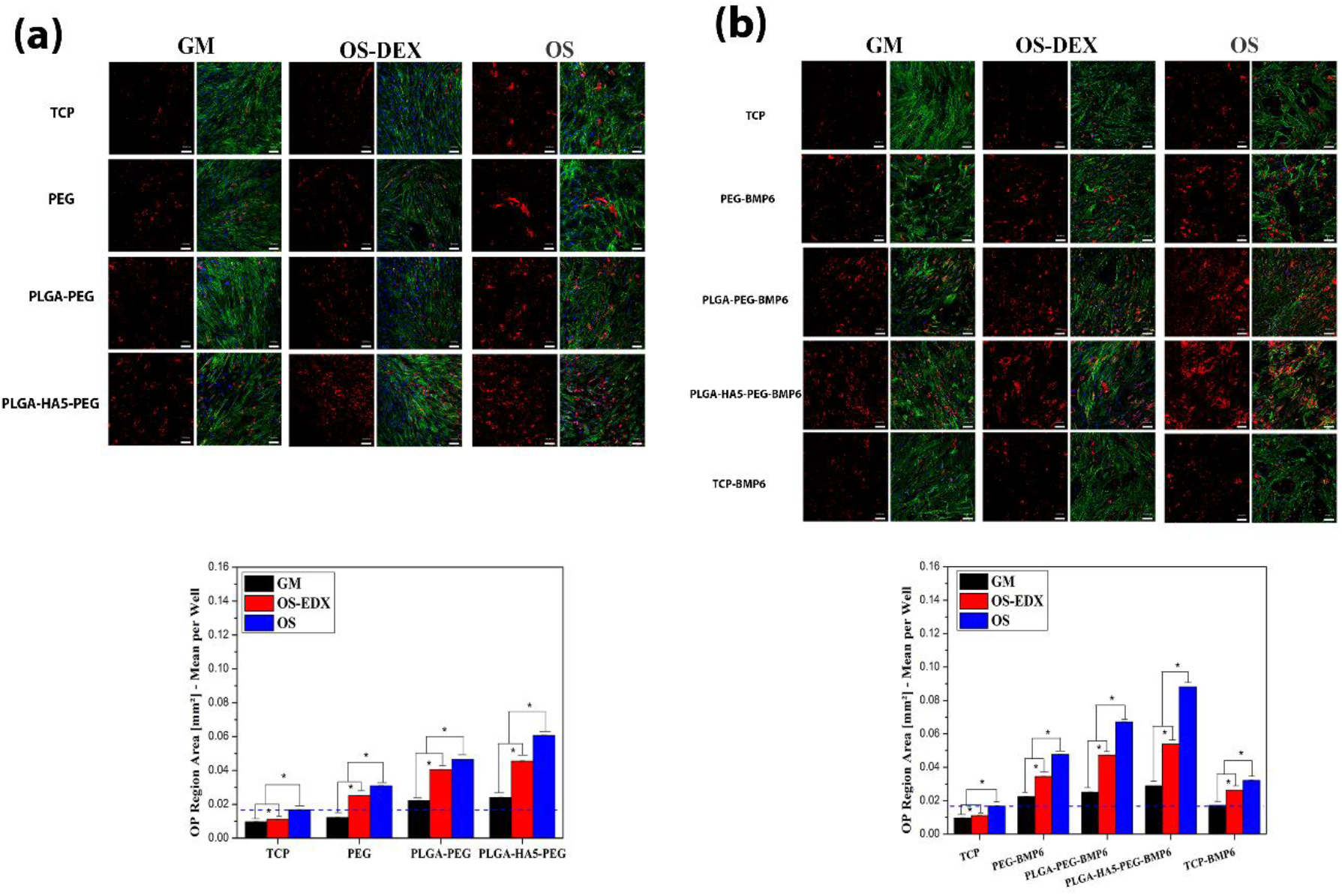
Immunofluorescence staining of OP in samples cultured with PEG hydrogels (a) before and (b) after BMP6 conjugation. OP expression was quantified based on immunofluorescence staining of samples cultured in GM, OS-DEX and OS. Scale bars, 200 μm; *P<0.05

**Figure 7.**
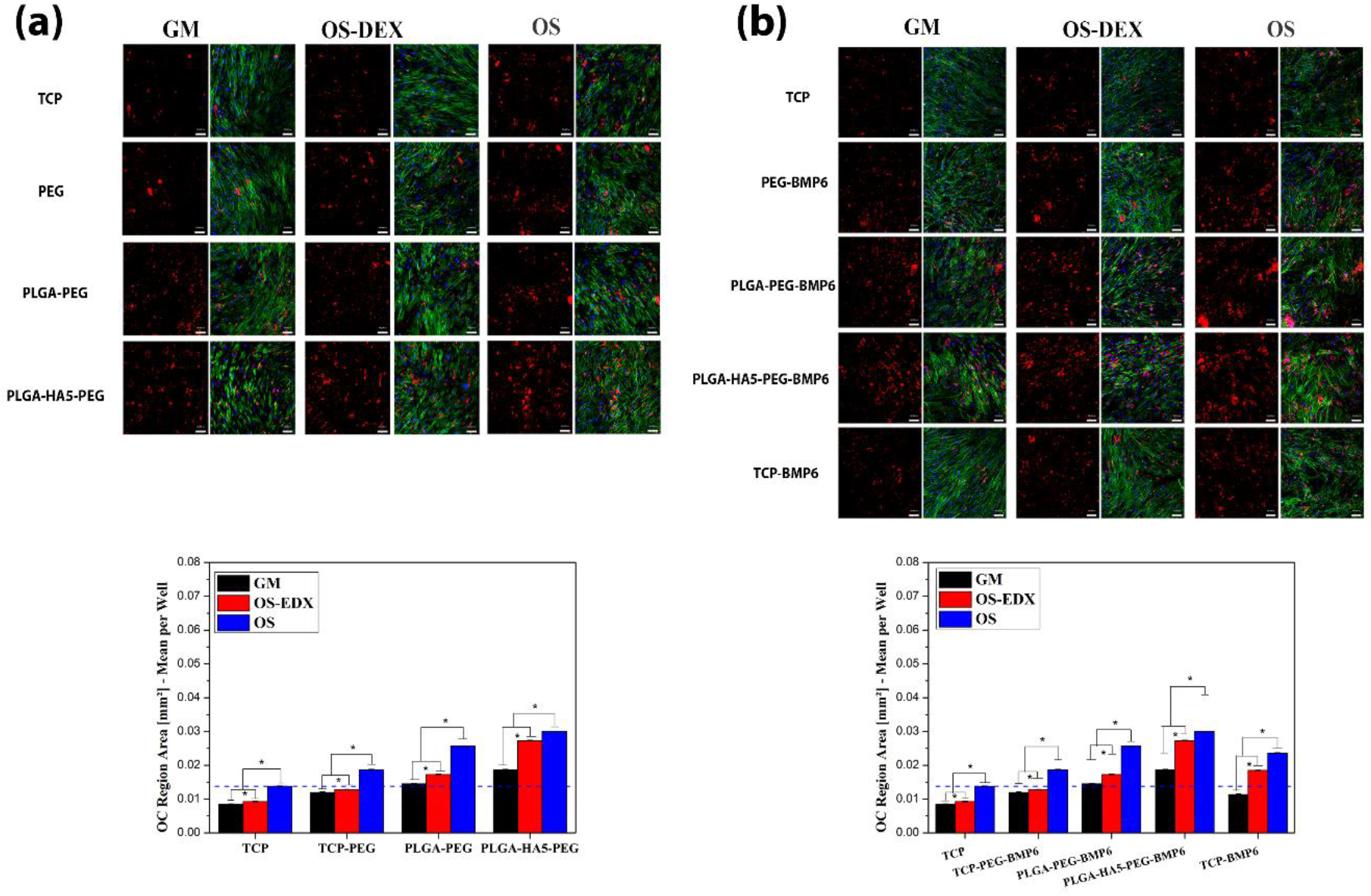
Immunofluorescence staining of OC in samples cultured with PEG hydrogels (a) before and (b) after BMP6 conjugation. OC expression was quantified based on immunofluorescence staining of samples cultured in GM, OS-DEX and OS. Scale bars, 200 μm; *P<0.05.

A comprehensive range of markers of osteogenic differentiation of hBMSCs, when cultured on bilayered scaffolds were assessed by qPCR. The expression of early osteogenic markers, including Runx2, ALP and Col1, as well as late osteogenic markers, including OP and OC, was investigated. Runx2 is known as the key regulatory gene for differentiation of mesenchymal progenitor cells to osteoblastic lineages via direct stimulation of downstream target genes including Col1, OP and OC ^[28]^. **Figure S13** shows that the incorporation of BMP6 conjugated PEG hydrogels into the scaffolds led to significantly higher Runx2 expression in all medium conditions compared to the TCP samples in similar medium conditions after 7 days. Notably, slightly higher Runx2 levels were observed after BMP6 conjugation into PEG hydrogels compared to the samples cultured with free BMP6 peptide in the same medium. However, considering the number of medium changes in the 7-day exposure period, these cells were exposed to almost 3 times more BMP6 peptide compared to BMP6 conjugated PEG samples. Although it is reported that DEX can contribute to osteogenic differentiation of hBMSCs by increasing Runx2 expression through the activation of WNT/β-catenin, TAZ and MKP-1 signals ^[29]^, the PLGA-HA5-PEG-BMP6 bilayered scaffolds in OS-DEX medium expressed a substantially higher level of Runx2 as compared to TCP in complete OS medium. This may be attributed to extensive phosphorylation of Smad 1/5/8 by the BMP6 conjugated PEG hydrogels that are known to recruit Runx2 to regulate the transcription of the osteogenesis ^[28b, 30]^. After 21 days, a similar trend was observed, regardless of medium conditions, but with significantly less Runx2 being expressed, indicating osteoblast maturation ^[31]^.

Next, Col1 **(Figure S14)** and ALP (**Figure S15)** gene expression was evaluated over the same timepoints. Extensive upregulation of both Col1 and ALP was observed when hBMSCs were exposed to BMP6 peptide. The Col1 and ALP expression on PLGA-HA5-PEG-BMP6 scaffolds cultured in OS-DEX medium were 12 fold and 7 fold higher, relative to TCP samples in complete OS medium. ALP and Col1 genes, were significantly upregulated in samples treated with BMP6 peptide in complete OS medium, highlighting the synergistic effect of BMP6 peptide and DEX to promote the osteogenic differentiation of hBMSCs. This is in line with previous research that demonstrated that the treatment of mouse C3H10T1/2 pluripotent stem cells with a combination of both BMP2 protein and DEX significantly upregulated both ALP and Col1 genes, as compared to treatment with each supplement alone _[32]_.

Osteoblastic maturation of hBMSCs cultured on bilayered scaffolds was further assessed by investigating changes in of late osteoblast-specific genes, OP (**Figure S16**) and OC (**Figure S17**). At day 21, there was significant upregulation of OP and OC for PLGA-HA5-PEG-BMP6 bilayered scaffolds, as compared to TCP samples in all medium conditions. The level of OP and OC expression for bilayered scaffolds cultured in OS medium was 55 fold and 35 fold, respectively, higher than that of TCP cultured in similar medium condition. Interestingly, even in medium lacking DEX, the PLGA-HA5-PEG-BMP6 expressed significantly higher levels of both genes compared to TCP in complete OS medium, indicating the osteogenic maturation of hBMSCs seeded in bilayered scaffolds, even in the absence of DEX or any osteogenic supplements. This finding supports previous research that reported BMP6 protein enhances gene expression level of Osterix, a transcription factor that regulates the expression of ECM proteins, such as osteopontin (OPN) in human dental follicle cells (hDFCs) cultured with in osteogenic induction medium without dexamethasone DEX ^[33]^.

### 2.3. Osteocytogenesis of hBMSCs cultured on osteon-like scaffolds from bilayered constructs

To recreate the 3D microenvironment seen by bone cells in the osteoid, osteon-like scaffolds (∼ 1mm diam., slightly larger than a native osteon (ave. ∼ 0.3 mm) but of the same order of magnitude) with concentric layers of hBMC-seeded PLGA-HA electrospun mats covered with BMP6p-conjugated PEG hydrogel were formed by simply rolling up our bilayered constructs. Furthermore, to assess whether physiologically-relevant stimuli would further promote osteocytogenesis in these osteon-like constructs, we next designed and fabricated a ‘plug-n-play’ prototype bioreactor system (**Figure S18a**) that allowed us to layer several osteon-like scaffolds whilst also exposing MSCs in concentric layers to maintenance or osteogenic medium for 21 days. To investigate the architecture of osteon-like scaffolds cultured in the bioreactor for 21 days, Micro CT analysis was performed. The micro CT images of acellular bilayered scaffolds incubated in GM (**Figure S18b**) and OS media (**Figure S18c**) showed a stable osteon-like architecture, maintaining the initial shape of the lumen structure surrounded by multilayers of PLGA-HA5-PEG-BMP6 matrix. This further confirmed that using this methodology no specialised equipment and processing conditions are necessary for the fabrication of multilayered tubular structures. Compared to more complex methods such as 3D printing or stress-induced rolling membrane (SIRM)^[34]^, tissue roll for analysis of cellular environment and response in 3D environment ^[35]^, by examining the 2D strip after unrolling, the need for complex equipment and expensive materials is eliminated.

In-vitro assessments of osteon-like scaffolds cultured in this bioreactor system were carried out after 21 days of incubation in GM and OS medium (**Figure 8**). Both scaffolds were cultured in GM and OS medium, and maintained their multilayered tubular structure, with multilayers of cell-laden sections (actin labelled ‘green’) of the scaffold forming the outer layers of the tubes, resembling osteon lamellae.

**Figure 8.**
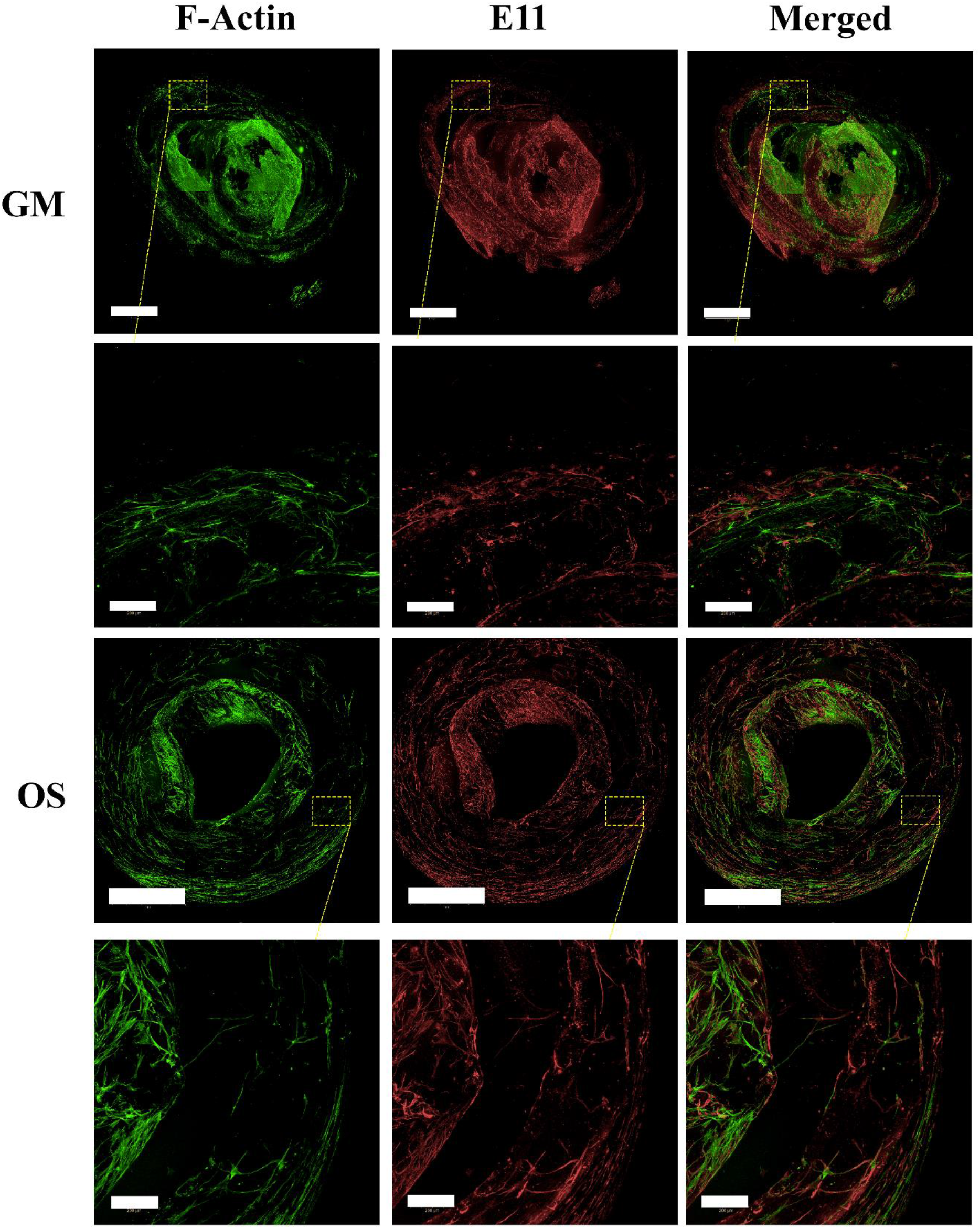
After 21 days, the osteon-like scaffolds were stained for E11 osteocyte markers. The osteocyte-like cells with multi-dendritic mulphology that stained positive for E11 was observed for both scaffolds were cultured in GM and OS.

Consequently, we analysed the E11 expression in osteon-like scaffolds to investigate the potential for osteocyte formation. E11 positive cells were observed throughout, for both osteon-like scaffolds cultured in GM or OS media. Enlarged images of E11 positive cells highlights the multi-dendritic osteocyte morphology with long processes, projecting from cell bodies with a high degree of connectivity between neighbouring cells, suggesting that the osteon-mimetic environment created by our system can promote osteocyte formation in the media lacking DEX or any exogenous osteogenic supplements after 21days.

We next conducted immunofluorescent staining for osteocyte markers in cross-sections of the osteon-like scaffolds after 21 days. Interestingly, both osteon-like scaffolds cultured in GM and OS stained positive for dentin matrix protein 1 (DMP1) (**Figure S19**), a regulatory gene for osteocyte maturation and phosphate metabolism ^[36]^. These images further confirmed that an integrated layer of cells forms along the surfaces of the neighbouring osteon-like lamellae of the scaffolds, showing significant potential for cell-cell communication across the multiple cultured osteon-like scaffolds in these bioreactors. The higher magnification images of osteon-like scaffolds shows (**Figure S19**) cells with multiple branching dendrites, stained positive for both E11 and DMP1 markers, projecting from cell bodies with a high degree of connectivity between neighbouring cells and even penetration of cells into the neighbouring layers of the scaffold, resembling the osteocyte lacuno-canalicular network in native cortical bone.

Since DMP1 is acidic protein that presents in the gap region between collagen type 1 fibrils during mineralization, regulating calcium binding and hydroxyapatite formation ^[37]^, we investigated the apatite formation of these osteocyte cells on sections of osteon-like scaffolds (**Figure S20**). Osteo-Image results demonstrate that, although some hydroxyapatite was formed on the surface of scaffolds, hydroxyapatite followed the E11 and DMP1 localisation, forming the interconnected networks. It has been shown previously that DMP1 can nucleate the formation of hydroxyapatite *in vitro* via a multistep process that begins with DMP1 binding calcium ions and initiating mineral deposition ^[38]^. This report has been confirmed further by He et al. ^[39]^ who reported that the negatively charged DMP1 fibril forms a stable structure by binding with calcium phosphate nanoparticles and protects the nascent mineral nuclei from further growth and precipitation. The observations throughout our scaffolds can thus be attributed to the nucleation of mineral at specific sites by DMP1, preventing spontaneous calcium phosphate precipitation in areas in which mineralization is not desirable.

## 3. Conclusion

A novel modular tissue engineering approach for the fabrication of a bilayered scaffold, composed of aligned PLGA-HA nanofibers and a BMP6 peptide-conjugated PEG-Nor/PEG-TH hydrogel, was described and validated. Firstly, the synthesised BMP6 peptides were shown to induce P-Smad 1/5/6 expression at even higher levels than the full length of the BMP6 protein. Secondly, the conjugation of the BMP6 peptide at low dosage to the PEG hydrogel enabled their sustained bioactivity for up to 21 days of hBMSC culture, along with their support of osteogenic differentiation by driving higher levels of both early and late osteogenic genes compared to full length BMP6 protein. Thirdly, the incorporation of the BMP6-functionalised hydrogels into bilayered scaffolds was shown to substantially increase ALP activity and calcium deposition of hBMSCs, as compared to TCP samples under all medium conditions. Moreover, the expression of major bone-specific transcription factors, including Runx2, Col1, OP and OC, on these hBMSC-seeded bilayered scaffolds were significantly higher in medium lacking DEX or any osteogenic supplements compared to TCP samples in complete OS medium. Finally, the maturation assessment of differentiated hBMSCs within the osteon-like scaffolds in our modular bioreactors confirmed the formation of osteocyte-like cells with dendritic processes similar to native osteocytes, secreting E11 and DMP1 osteocyte markers, even in medium lacking any osteogenic supplements after 21 days of culture. This facile ex vivo culture model thus provides a novel production platform for osteocyte-like cells from human MSCs and the creation of ‘plug-n-play’ cortical bone mimics *ex vivo*.

## 4. Experimental Section

### 4.1. Materials

PLGA (LA/GA = 75/25, Inherent Viscosity 0.55–.75 Dl/g in CHCL_3_) was purchased from LACTEL USA. Calcium nitrate and ammonium phosphate dibasic were purchased from Sigma Aldrich (Australia). 8-arm PEG-NH_2_ (10 kDa) (Jenkem) and 4-arm PEG-TH (10 kDa) were purchased from JenKem. Cysteamine was provided by Alfa Aessar. Human bone marrow mesenchymal stem cells (Lonza), low glucose DMEM and FBP (Gibco, 10567-022), All other chemicals were purchased from Sigma Aldrich (Australia), if not otherwise mentioned.

### 4.2. Maintenance and Expansion of hBMSCs

The hBMSCs (Donors 4287, 7219 and 8006, Lonza, Switzerland) at passage 2 and expanded in growth medium (GM), which was α Minimum Essential Medium (α-MEM, GlutaMAX Supplement, no nucleosides, TFS) supplemented with 10% fetal bovine serum (FBS), penicillin (100 units/mL) and streptomycin (100 μg/mL) (P/S) (Gibco, Australia), 1% sodium pyruvate and 0.25% ascorbic acid. The cells were cultured under standard conditions (humidified, 37 ºC, 95% air, 5% CO_2_). Media was changed every second day until 80% confluence. hBMSCs were subcultured using TrypLE™ Express (Life Technologies, CA, USA) and replated at 4000 cells.cm^−2^. All the experiments were performed at passage 5.

### 4.3. Peptide Synthesis, Purification and Characterisation

The BMP6-peptide mimics the receptor-binding domain of the BMP-6 protein synthesised by using Fmoc chemistry at the 0.2 mmol scale. 20% piperidine in *N, N*′-dimethylformamide (DMF) was used for deprotection of the Fmoc group. Subsequent Fmoc protected amino acids were coupled to the free amino group using *O*-(benzotriazole-1-yl)-*N, N, N*′, *N*′-tetramethyluronium hexafluorophosphate (HBTU) as the coupling reagent. After each deprotection and coupling step, the resin was washed with DMF (∼5 mL) 6 times for 3 min. After coupling the last amino acid, excess reagents were removed by washing 6 times in DMF for 5 min, followed by five washing steps using DCM for 2 min (∼5 mL). The resin was then vacuum dried for 2 hours prior to cleavage. Peptides were cleaved from the resin and deprotected in trifluoroacetic acid (TFA) with 2.5% deionised water and 1.25% dithiothreitol (DTT, Research Products International) and 1% triisopropylsilane (TIPS) under nitrogen flow for 2 hours. Peptides were then precipitated in 20 mL per gram of resin of ice-cold diethyl ether and centrifuged at 4 °C at 4400 rpm for 5 min. The residue was washed with diethyl ether before resuspension in deionised water. The solution was frozen with liquid nitrogen and lyophilised overnight to yield a fluffy white powder.

The base amino acid sequences for the BMP6 peptide were as per reported (YVPLPCCAPTKLNAISVLYF) ^[11a]^. Then we employed thiol-end click chemistry, adding a cysteine group to the N-terminus of the peptide to conjugate our BMP6 peptide to a polyethylene glycol (PEG)-star polymer. The chemical composition and the molecular structure of the synthesised cysteine-terminated BMP-6 peptide are shown in **Figure S21**. Synthesised peptides were then purified and characterised using high-pressure liquid chromatography (HPLC) (**Figure S22**) and matrix-assisted laser desorption ionisation (MALDI) mass-spectroscopy (**Figure S23**). The integration of peaks from the HPLC result for the BMP6 peptide showed the peptide purity. The presence of impurities in peptides can lower or alter their binding capability ^[25, 40]^; therefore, a high-quality peptide product (with greater than 90% purity) is required for accurate results in binding assays and *in vitro* studies.

### 4.4. High-Performance Liquid Chromatography

High-performance liquid chromatography (HPLC) (Agilent Technologies 1260 Infinity) in an acetonitrile/water gradient under acidic conditions on a Phenomenex Luna 5um C18 (2) column (5 μm pore size, 100 Å particle size, 150 × 4.6 mm) was used to check the presence of the main peak representing peptide and other components (impurities). Preparative HPLC with Phenomenex Luna 10 μm C18 (2) column (10 μm pore size, 100 Å particle size, 150 × 21.2 mm) column was used to separate peaks and purify the peptide.

### 4.5. Matrix-Assisted Laser Desorption/Ionisation Time-Of-Flight Mass Spectrometry

Matrix-Assisted Laser Desorption/Ionisation Time-Of-Flight Mass Spectrometry (MALDI-TOF MS) measurement was performed on a Bruker Autoflex III MALDI-TOF mass spectrometer. The purified peptide was dissolved in water at 1 mg ml^−1^. A stock solution of the matrix was prepared by dissolving 10 mg of HCCA-ALPHA-cyano-4-hydroxycinnamic acid in 1 mL of water. The sample and matrix solutions were combined in the ratio of 100 to 10 μl, respectively. A 1 μl aliquot of the mixture was then applied to the Prespotted Anchor Chip MALDI plate and air-dried at ambient temperature (20 °C). The measurement was performed at an acceleration voltage of 19 kV in positive ion reflection mode. Suitable values for laser power, gain, and shots were determined for each sample to produce the best quality data (best resolution, reduced fragmentation, and ion statistics).

### 4.6. Synthesis and Characterisation of PEG-based Hydrogels

#### 4.6.1. PEG Functionalisation

Norbornene-functionalised PEG was synthesised (PEG/Nor) as described elsewhere ^[41]^. Briefly, the 8-arm PEG-NH_2_ (*M*_w_ 10 000) was dissolved in dichloromethane (DCM). In a separate flask, norbornene carboxylic acid (5 Equiv.) was activated by HBTU and HOBT (4 Equiv.) in DMF for 3 min at room temperature. DIEA (6 Equiv.) was added to the activated acid solution and further reacted for 5 min under nitrogen. The mixture was then added dropwise to the flask containing 8-arm PEG-NH_2_ and allowed to react overnight under nitrogen. Afterwards, the reaction was filtered and concentrated under rotary evaporation, followed by precipitation in ice-cold diethyl ether. The filtered PEG-Nor powder was then dissolved in water and lyophilised. PEG-Nor was dialysed against the water with a molecular cut-off of 5 kDa for 3 days before being used. The degree of functionalisation (>85%) was measured using proton Nuclear magnetic resonance spectroscopy (NMR).

#### 4.6.2. PEG/Nor-PEG/TH Hydrogel Formation

8-arm norbornene functionalised PEG was cross-linked with 4-arm PEG-thiol in the presence of lithium phenyl-2,4,6-trimethylbenzoylphosphinate (LAP) as a photoinitiator under UV light (intensity of 180 mW/cm^−1^) for 10 seconds at room temperature. A stock solution of 30% (w/v) PEG-norbornene and 10% (w/v) PEG-thiol and 5 mM LAP was prepared. The thiol to norbornene ratio was 1:1. LAP was added to a final concentration of 0.08% (w/v). PBS buffer was used to make up the solution to the final desired PEG-norbornene concentration. For BMP6 peptide conjugated gels, peptides were added to the gel solution to make up different concentrations (0.1, 0.5, 1, 2 and 10 μg/ml). The concentration of LAP was kept at 0.05% (w/v).

#### 4.6.3. Nuclear magnetic resonance spectroscopy

Quantitative ^1^H NMR (300 or 500 MHz) spectra for PEG-norbornene were acquired on a Bruker NMR 300 Hz Ultrasheild ((Bruker Chemical Analysis, PL, USA). Samples for NMR analysis were prepared by dissolution of 10 mg of product in D_2_O or CDCl_3_ on a glass vial (Sigma-Aldrich). After complete dissolution, samples were transferred to NMR tubes. The degree of substitution of norbornene on PEG macromers was calculated from the integral of the methylene resonance from PEG (δ = 3.7 ppm) and the double bond from norbornene (δ = 6.0 and 5.9 ppm), normalising the number of contributing protons. Integration of the peaks for PEG and norbornene confirmed that the degree of functionalisation of the PEG with norbornene was 88%. Data analysis was performed using Mnova NMR software (Mestrelan Research, Galicia, Spain).

### 4.7. Cryo-SEM Imaging

The samples for cryo-SEM were prepared using the high-pressure freezing (HPF) method as described before ^[42]^. The samples were then transferred under vacuum to the cold (<100 K) pre-chamber of the Gatan Alto 2500 cryo-preparation and coating station. Inside the cryo-preparation chamber, samples were fractured with a Gatan precision cold rotary fracture knife (ALT 335) to provide a clean surface uncontaminated by atmospheric water. Samples were routinely sublimed at −105 °C, sputter-coated in the cryo-preparation chamber with Pt for 240 seconds at 10 mA (circa 5–10 nm thickness), then transferred to the Polaron LT7400 cryo prep chamber that was attached to a Philips XL30 field emission scanning electron microscope (FESEM) for imaging. The samples were sublimed (within the cryo prep chamber) for 45 min at −80 °C before being placed into the SEM chamber at −190 °C and imaged using an accelerating voltage of 2 kV.

### 4.8. Rheometry

In situ dynamic rheometry was performed with parallel-plate geometry on an ARG2 rheometer (TA Instruments). UV light at 365 nm (10 mW cm^2^) was directed through a flat quartz plate through the sample, while time sweep measurements were made at a 100 μm gap, 10 Pa stress and 10 rad s. The gel point was taken as the storage/loss moduli crossover point, which indicates a conversion in the vicinity of the precise gel point.

### 4.9. Preparation of the nHA and Nanofibrous Scaffold

The rod shape nHA was synthesised based on a previously published method from our group ^[43]^. Briefly, a 50 mL aliquot of 0.5 mol L^−1^ Ca(NO_3_)_2_ was heated to 70 °C, and the same amount of 0.3 mol L^−1^ (NH_4_)2HPO_4_ was added dropwise for 1 hour while stirring and maintaining the pH above 9 and the temperature at 70 °C. The resulting solution was further allowed to stir for one hour at the same temperature. The formed nHA was then washed by centrifugation at 1000 rpm for 5 minutes, exchanging the supernatant and resuspending the particles several times using Milli-Q water until a neutral pH of the washing solution was obtained. It was then dried in an oven at 50 °C overnight before powdering.

PLGA nanofibers and PLGA-nHA nanocomposites were fabricated using an electrospinning technique based on a previously published method from our group ^[44]^. First, PLGA was dissolved in CHCL_3_/DMF (7/3 v/v) with a 20% (w/v) concentration. The mixture was stirred vigorously for 2 hours at room temperature before the spinning. For fabrication of PLGA-HA nanocomposites, different amounts of nHA (0, 1, 5, 10, 15 and 20 wt% based on PLGA content) were uniformly dispersed in DMF for 20 minutes under sonication. Then half of the PLGA was dissolved in DMF, and the nHA was mixed into this solution under magnetic stirring for 1h. Finally, the rest of PLGA was dissolved in the solution, and it was stirred for 1 more hour. The solution was ejected from a 5 mL syringe with a steel needle (inner diameter: 12 mm) using a programmable syringe pump (Top 5300, Japan) at a rate of 0.7 mL/h. A high voltage of 18 kV was applied to the solutions while aligned nanofibers were collected between two copper blade arms.

### 4.10. Characterisation of the nHA, PLGA and PLGA-HA Nanocomposites

The surface morphology of the scaffolds was examined by scanning electron microscopy (SEM HITACHI SU3500) with energy-dispersive X-ray spectroscopy for elemental analysis. The specimens were sputtered with a thin layer of iridium to reduce the surface charging of the samples. The SEM was operated at 15–20 kV to observe the surface features of the scaffolds. Transmission electron microscopy (JEOL JEM-1010 TEM) was used to observe the morphology of nHA and its dispersion into the matrices.

### 4.11. Cell Culture and Osteogenic Differentiation

Before cell culture on PLGA and PLGA-HA substrates, they were placed on a tissue culture plate and incubated for 15 min in 70 vol % ethanol and rinsed with phosphate buffer saline solution (PBS tablets, Thermo-Fisher, Australia) three times and once with low α-MEM, after which they were placed in new tissue culture plates. Then, hBMSCs were seeded on the surface of the scaffolds at 4 × 10^4^ cells/cm^2^ for 24 hours under standard cell culture conditions. After the cell attachment, the surface of cell-seeded scaffolds was covered with a layer of either non-conjugated or conjugated BMP6 peptide PEG-based hydrogels to fabricate the bilayered scaffolds. The culture media was then changed to either the complete osteogenic differentiation media (OS) (α-MEM, 10% fetal bovine serum, 1% penicillin-streptomycin, 100 nM Dexamethasone + 100 μM Ascorbic acid 2-phosphate + 10 mM β-glycerolphosphate) or the OS medium lacking Dexamethasone (OS-DEX). The medium was changed every 2–3 days during 7, 14 and 21 days of *in vitro* culture.

### 4.12. Cell Viability

Cell seeding on sterilised PLGA-based substrates was performed at a density of 10000 cells/cm^2^ and in FBS free medium (α-MEM and 1% penicillin-streptomycin). Qualitative evaluation of the presence of viable cells was accomplished using a Live/Dead viability kit (Thermo Fisher Scientific L7013) after culture for 24 hours. Samples were incubated with the staining solution at 37 °C for 30 min in a 24-well plate, and the cells were imaged under an Operetta fluorescence microscope (Operetta, Perkin Elmer).

### 4.13. Morphological Observation of hBMSCs on Nanocomposite Fibres

After 21 days of hBMSC differentiation, cell morphology was observed by SEM with an accelerating voltage of 15–20 kV. The hBMSCs cultured on nanofibres were washed with PBS, fixed with 2.5% glutaraldehyde and then rinsed with water. The samples were dehydrated through a series of graded alcohol solutions and then air-dried overnight. Before observation, the cells were coated with iridium using a Q150TS Quorum coater.

### 4.14. Immunofluorescence Staining

At the predetermined time points, the cultured cells were washed and subsequently fixed with 4% paraformaldehyde for 10 min, followed by cell permeabilisation and blocking of non-specific protein-protein interaction using a 1% bovine serum albumin (BSA)/10% normal goat serum in solution 0.1% (w/v) Triton 100-X solution for 5 min. After PBS rinsing, samples were blocked in a 0.1 PBS-Tween for 1 hour. The blocked samples were incubated with Phospho-Smad 1/Smad5/Smad9 (D5B10) Rabbit mAb (Cell Signalling Technology 13820), osteoblast-specific marker protein primary antibody Collagen I (Col1) (Anti-Collagen I antibody ab34710), osteopontin primary antibody (OP) (Anti-Osteopontin antibody ab8448) and osteocalcin primary antibody (OC) overnight at 4 °C. Following incubation with the primary antibody, the samples were washed and incubated with a secondary antibody: Goat Anti-Rabbit IgG H&L (Alexa Fluor® 568, Life Technologies, Australia) for Col I, Goat Anti-Rabbit IgG H&L (Alexa Fluor® 647, Life Technologies, Australia) for OP and Donkey Anti-Rabbit for OC. Subsequently, Phalloidin 488 (Life Technologies, Australia) and Hoechst 33342 were used to stain F-actins and nuclei, respectively. The cells were visualised via LSM Zeiss 710 confocal microscope or the confocal Operetta (Operetta, Perkin Elmer). For immune staining of osteocyte markers, primary antibodies against E11 (1:1000 ab10288, Abcam) and DMP1 (1:1000 ab103203, Abcam) were diluted using the antibody dilution buffer (1% BSA + 0.05% Tween-20). Following overnight incubation with the primary antibody, the samples were washed and incubated with secondary antibodies for 1 hour at room temperature.

### 4.15. Alkaline Phosphatase Activity Staining

To characterise the osteogenic differentiation of hBMSCs on the bilayered scaffolds, the samples were stained for alkaline phosphatase (ALP). After 7 days in the respective culture conditions, the samples were washed with PBS and stained with a solution containing Fast Blue B and naphthol AS-MX ALP solution. The samples were eventually fixed using 4% formaldehyde.

### 4.16. Alizarin Red Staining

Alizarin red staining was performed to evaluate the amount of ECM produced by hBMSCs qualitatively and quantitatively. After 21 days of culture, the scaffolds were fixed in 4% formaldehyde for 10 min. Then, the scaffolds were washed with distilled water and stained with ARS (0.1%) for 30 min at room temperature. After washing several times with distilled water, the scaffolds were examined under an upright optical microscope (IX51 Inverted Microscope, Olympus, Japan). The deposited calcium was then dissolved by 10% cetylpyridinium chloride monohydrate (Sigma-Aldrich, U.S.A.) and quantified using a microplate reader at 540 nm wavelength. Acellular scaffolds in water or medium were analysed as the control to rule out the effects of n-HA on calcium deposition outcomes.

### 4.17. Osteo-Image Staining

Osteo-Image (Lonza, Walkersville, USA) was used as a specific stain for hydroxyapatite (HA). The samples were washed once with PBS and subsequently fixed with 4% formaldehyde for 10 min, followed by washing twice with Osteo-Image wash buffer (diluted 1:10 in deionised water). An appropriate amount of diluted staining reagent (diluted 1:100 in staining reagent dilution buffer) was added to each sample, and the mixture was incubated for 30 min, protected from light. Then, the staining reagent was removed, and the samples were washed three times with the wash buffer. The samples were analysed under a confocal microscope.

### 4.18. Quantitative Real-Time Reverse Transcription Polymerase Chain Reaction

After 7, 14 or 21 days of differentiation on PLGA-based substrates, total RNA was extracted using NucleoSpin RNA Set for NucleoZOL (Macherey-Nagel) according to the manufacturer’s protocol. cDNA was generated from 200 ng of RNA using SuperScript III First-Strand Synthesis SuperMix (Invitrogen, Life Technologies). The cDNA was stored at −20 °C until further analyses. RT-PCRs were prepared on a total volume of 10 μL with 5 μL SYBR green (Thermo-Fisher), 0.2 μM forward and reverse primers, and 100 ng cDNA and DEPC treated water. For no-RT controls, an equivalent volume of DNase and RNase-free water (Sigma) was used in place of RT Enzyme Mix. A CFX96™ IVD Real-Time PCR system (BioRad) was used with a thermal cycle of 50 °C for 2 min, 95 °C for 2 min, and then 95 °C for 15 s and 60 °C for 30 s for a total of 40 cycles. Ct values of RT-PCR were normalised against the housekeeping gene and analysed using the ΔΔCt model [47]. The expression of bone-associated genes, including Smad 1/5/8, ALP, Col1, Runt-related transcription factor 2 (Runx 2) and osteopontin (OP) and osteocalcin (OC) was confirmed via quantitative RT-PCR. The primer sequences used for RT-PCR were Smad 1 forward, GGCGGCATATTGGAAAAGGAGTT; Smad 1 reverse, GTTGCAGTTCCGACTTTGCACA. Smad 5, GAGTGGAGAGTCCAGTCTTACCTC; Smad 5, TTGTGGCTCAGGTTCCTAAACTGAAC. Smad 9 forward, AGACATATAGGAAAGGGTGTGCACTT; Smad 9 reverse, TGCAGTTCCGGCTCTGCA Runx2 forward, AGTGATTTAGGGCGCATTCCT; Runx2 reverse, GGAGGGCCGTGGGTTCT. ALP forward, GGGAACGAGGTCACCTCCAT; ALP reverse, TGGTCACAATGCCCACAGAT. Col1 forward, CCTGCGTGTACCCCSCTCA; Col1 reverse, ACCAGACATGCCTCTTGTCCTT. OP forward, ACCTGAACGCGCCTTCTG; OP reverse, CATCCAGCTGACTCGTTTCATAA. OC forward; AGCAAAGGTGCAGCCTTTGT, OC reverse; GCGCCTGGGTCTCTT GAPDH forward, ATGGGGAAGGTGAAGGTCG; GAPDH reverse, TAAAAGCAGCCCTGGACC.

### 4.19. Microcomputed Tomography (Micro CT)

Micro CT analysis was carried out using SkyScan 1172 high-resolution micro-CT imaging system. Micro CT images were taken at a resolution of 18 μm with no filter, under a voltage of 100 kV and a current of 100 μA. 3D models of the samples were created as needed using CTVol software provided by Skyscan.

### 4.20. Bioreactor Fabrication

Bioprinting manufacturing technology is employed using the SLA bioprinter (Formlabs Form 2 Starter Pack) to print the customised bioreactor device. High Temp Resin (heat deflection temperature (HDT) of 238 °C @ 0.45 MPa) was used. The printed constructs were post-cured with a 405 nm light source for 60 minutes.

### 4.21. Statistical Analysis

Statistical significance was calculated by one-way ANOVA to assess the statistical significance (p < 0.05) of the differences between results. For differentiation experiments, analysis of ALP activity assay, ALP, Alizarin Red and Osteo-Image staining was quantified from a minimum of 12 images recorded from 3 samples (triplicates), repeated with three donors. Three biological replicates were performed for all other cellular experiments, and the data were pooled together (n ≥ 9).

## Supporting Information

Supporting Information is available from the Wiley Online Library or from the author.

## Acknowledgements

This work was supported by the Australian Research Council Discovery Grants Scheme (DP190101969). This work was performed in part at the Queensland node of the Australian National Fabrication Facility, a company established under the National Collaborative Research Infrastructure Strategy to provide nano and microfabrication facilities for Australia’s researchers.

## Data Availability Statement

Research data are not shared.

## References

[1] N. Lohse, N. Moser, S. Backhaus, T. Annen, M. Epple, H. Schliephake, Journal of Controlled Release 2015, 220, 201.

[2] M. Wu, G. Chen, Y.-P. Li, Bone Research 2016, 4, 16009.

[3] a) C. Laurencin, Y. Khan, S. F. El-Amin, Expert Rev. Med. Devices 2006, 3, 49; b) E. Segredo-Morales, P. García-García, C. Évora, A. Delgado, Journal of Drug Delivery Science and Technology 2017, 42, 107.

[4] a) T. J. Blokhuis, J. J. C. Arts, Injury 2011, 42, S26; b) P. Gentile, A. M. Ferreira, J. T. Callaghan, C. A. Miller, J. Atkinson, C. Freeman, P. V. Hatton, Advanced Healthcare Materials 2017, 6, 1601182.

[5] K. Alvarez, H. Nakajima, Materials 2009, 2, 790.

[6] a) M. Zhang, R. Lin, X. Wang, J. Xue, C. Deng, C. Feng, H. Zhuang, J. Ma, C. Qin, L. Wan, J. Chang, C. Wu, Science Advances 2020, 6, eaaz6725; b) D. Barati, O. Karaman, S. Moeinzadeh, S. Kader, E. Jabbari, Regenerative biomaterials 2019, 6, 89; c) X. Chen, A. Ergun, H. Gevgilili, S. Ozkan, D. M. Kalyon, H. Wang, Biomaterials 2013, 34, 8203.

[7] a) W. Cui, Q. Liu, L. Yang, K. Wang, T. Sun, Y. Ji, L. Liu, W. Yu, Y. Qu, J. Wang, Z. Zhao, J. Zhu, X. Guo, ACS Biomaterials Science & Engineering 2018, 4, 211; b) B. Huang, Y. Yuan, S. Ding, J. Li, J. Ren, B. Feng, T. Li, Y. Gu, C. Liu, Acta biomaterialia 2015, 27, 275; c) V. Rosen, Cytokine Growth Factor Rev. 2009, 20, 475; d) H. Nie, M.-L. Ho, C.-K. Wang, C.-H. Wang, Y.-C. Fu, Biomaterials 2009, 30, 892; e) B. Bragdon, O. Moseychuk, S. Saldanha, D. King, J. Julian, A. Nohe, Cell. Signal. 2011, 23, 609.

[8] H. K. Kim, J. S. Lee, J. H. Kim, J. K. Seon, K. S. Park, M. H. Jeong, T. R. Yoon, Experimental & Molecular Medicine 2017, 49, e328.

[9] a) H. H. Luu, W. X. Song, X. Luo, D. Manning, J. Luo, Z. L. Deng, K. A. Sharff, A. G. Montag, R. C. Haydon, T. C. He, J. Orthop. Res. 2007, 25, 665; b) H. Cheng, W. Jiang, F. M. Phillips, R. C. Haydon, Y. Peng, L. Zhou, H. H. Luu, N. An, B. Breyer, P. Vanichakarn, JBJS 2003, 85, 1544; c) A. L. Grab, D. Hose, P. Horn, E. A. Cavalcanti-Adam, A. Seckinger, M. Müller, Particle & Particle Systems Characterization 2021, 38, 2000263.

[10] a) A. E. Mercado, J. Ma, X. He, E. Jabbari, Journal of Controlled Release 2009, 140, 148; b) Y. Robinson, C. E. Heyde, S. K. Tschöke, M. A. Mont, T. M. Seyler, S. D. Ulrich, Expert Rev. Med. Devices 2008, 5, 75.

[11] a) Y. J. Choi, J. Y. Lee, C. P. Chung, Y. J. Park, Biomater Res 2014, 5, 19; b) O. Mizrahi, D. Sheyn, W. Tawackoli, I. Kallai, A. Oh, S. Su, X. Da, P. Zarrini, G. Cook-Wiens, D. Gazit, Z. Gazit, Gene Ther. 2012, 20, 370; c) F. Kugimiya, H. Kawaguchi, S. Kamekura, H. Chikuda, S. Ohba, F. Yano, N. Ogata, T. Katagiri, Y. Harada, Y. Azuma, J. Biol. Chem. 2005, 280, 35704; d) Y. Cao, Q. Tan, J. Li, J. Wang, Braz. J. Med. Biol. Res. 2020, 53.

[12] a) Y. Chen, F. Ding, H. Nie, A. W. Serohijos, S. Sharma, K. C. Wilcox, S. Yin, N. V. Dokholyan, Arch. Biochem. Biophys. 2008, 469, 4; b) C. Li, C. Vepari, H.-J. Jin, H. J. Kim, D. L. Kaplan, Biomaterials 2006, 27, 3115.

[13] a) X. Niu, Q. Feng, M. Wang, X. Guo, Q. Zheng, Journal of Controlled Release 2009, 134, 111; b) Y. J. Lee, J.-H. Lee, H.-J. Cho, H. K. Kim, T. R. Yoon, H. Shin, Biomaterials 2013, 34, 5059.

[14] a) Bo-Ram Kim, Thuy Ba Linh Nguyen, Young-Ki Min, B.-T. Lee, Tissue Engineering Part A 2014, 20, 3279; b) L. Li, G. Zhou, Y. Wang, G. Yang, S. Ding, S. Zhou, Biomaterials 2015, 37, 218.

[15] T. Andric, A. C. Sampson, J. W. Freeman, Materials Science and Engineering: C 2011, 31, 2.

[16] L. I. Plotkin, T. Bellido, Nature Reviews Endocrinology 2016, 12, 593.

[17] M. Soheilmoghaddam, H. Padmanabhan, J. J. Cooper-White, Biomaterials Science 2020, 8, 5677.

[18] J. E. Frith, D. J. Menzies, A. R. Cameron, P. Ghosh, D. L. Whitehead, S. Gronthos, A. C. W. Zannettino, J. J. Cooper-White, Biomaterials 2014, 35, 1150.

[19] a) B. L. Rosenzweig, T. Imamura, T. Okadome, G. N. Cox, H. Yamashita, P. ten Dijke, C. H. Heldin, K. Miyazono, Proc. Natl. Acad. Sci. U. S. A. 1995, 92, 7632; b) T. Nohno, T. Ishikawa, T. Saito, K. Hosokawa, S. Noji, D. H. Wolsing, J. S. Rosenbaum, J. Biol. Chem. 1995, 270, 22522.

[20] a) M. Beederman, J. D. Lamplot, G. Nan, J. Wang, X. Liu, L. Yin, R. Li, W. Shui, H. Zhang, S. H. Kim, Journal of biomedical science and engineering 2013, 6, 32; b) T. Ebisawa, K. Tada, I. Kitajima, K. Tojo, T. K. Sampath, M. Kawabata, K. Miyazono, T. Imamura, Journal of cell science 1999, 112, 3519.

[21] I. Simeoni, J. B. Gurdon, Dev. Biol. 2007, 308, 82.

[22] F. Chen, X. Lin, P. Xu, Z. Zhang, Y. Chen, C. Wang, J. Han, B. Zhao, M. Xiao, X.-H. Feng, Mol. Cell. Biol. 2015, 35, 1700.

[23] M. Yuasa, T. Yamada, T. Taniyama, T. Masaoka, W. Xuetao, T. Yoshii, M. Horie, H. Yasuda, T. Uemura, A. Okawa, PLoS One 2015, 10, e0116462.

[24] H. Follet, G. Boivin, C. Rumelhart, P. J. Meunier, Bone 2004, 34, 783.

[25] H. Hezaveh, S. Cosson, E. A. Otte, G. Su, B. D. Fairbanks, J. J. Cooper-White, Biomacromolecules 2018, 19, 721.

[26] M. Prideaux, C. Schutz, A. R. Wijenayaka, D. M. Findlay, D. G. Campbell, L. B. Solomon, G. J. Atkins, Bone 2016, 88, 64.

[27] G. Y. Rochefort, S. Pallu, C. L. Benhamou, Osteoporos. Int. 2010, 21, 1457.

[28] a) M. Mohammadi, M. Alibolandi, K. Abnous, Z. Salmasi, M. R. Jaafari, M. Ramezani, Nanomedicine: Nanotechnology, Biology and Medicine 2018, 14, 1987; b) R. L. Huang, Y. Yuan, J. Tu, G. M. Zou, Q. Li, Cell Death & Disease 2014, 5, e1187.

[29] F. Langenbach, J. Handschel, Stem Cell. Res. Ther. 2013, 4, 117.

[30] a) W.-G. Jang, E.-J. Kim, D.-K. Kim, H.-M. Ryoo, K.-B. Lee, S.-H. Kim, H.-S. Choi, J.-T. Koh, The Journal of biological chemistry 2012, 287, 905; b) M. Phimphilai, Z. Zhao, H. Boules, H. Roca, R. T. Franceschi, Journal of bone and mineral research : the official journal of the American Society for Bone and Mineral Research 2006, 21, 637.

[31] Z. Maruyama, C. A. Yoshida, T. Furuichi, N. Amizuka, M. Ito, R. Fukuyama, T. Miyazaki, H. Kitaura, K. Nakamura, T. Fujita, N. Kanatani, T. Moriishi, K. Yamana, W. Liu, H. Kawaguchi, K. Nakamura, T. Komori, Dev. Dyn. 2007, 236, 1876.

[32] Y. Mikami, M. Asano, M. J. Honda, M. Takagi, J. Cell. Physiol. 2010, 223, 123.

[33] K. Takahashi, N. Ogura, H. Aonuma, K. Ito, D. Ishigami, Y. Kamino, T. Kondoh, Arch. Oral Biol. 2013, 58, 690.

[34] Y. Jin, N. Wang, B. Yuan, J. Sun, M. Li, W. Zheng, W. Zhang, X. Jiang, Small 2013, 9, 2410.

[35] a) A. Bandyopadhyay, I. Mitra, S. Bose, Current osteoporosis reports 2020, 18, 505; b) D. Rodenhizer, E. Gaude, D. Cojocari, R. Mahadevan, C. Frezza, B. G. Wouters, A. P. McGuigan, Nature Materials 2016, 15, 227.

[36] Y. Lu, B. Yuan, C. Qin, Z. Cao, Y. Xie, S. L. Dallas, M. D. McKee, M. K. Drezner, L. F. Bonewald, J. Q. Feng, Journal of bone and mineral research : the official journal of the American Society for Bone and Mineral Research 2011, 26, 331.

[37] a) J. Shao, Y. Zhou, Y. Xiao, Bone 2018, 108, 165; b) M. B. Schaffler, O. D. Kennedy, Current osteoporosis reports 2012, 10, 118.

[38] G. He, T. Dahl, A. Veis, A. George, Nature Materials 2003, 2, 552.

[39] G. He, S. Gajjeraman, D. Schultz, D. Cookson, C. Qin, W. T. Butler, J. Hao, A. George, Biochemistry 2005, 44, 16140.

[40] L. Gil-Guerrero, J. Dotor, I. L. Huibregtse, N. Casares, A. B. López-Vázquez, F. Rudilla, J. I. Riezu-Boj, J. López-Sagaseta, J. Hermida, S. Van Deventer, J. Bezunartea, D. Llopiz, P. Sarobe, J. Prieto, F. Borrás-Cuesta, J. J. Lasarte, The Journal of Immunology 2008, 181, 126.

[41] A. Raza, C.-C. Lin, Macromol. Biosci. 2013, 13, 1048.

[42] a) R. Aston, K. Sewell, T. Klein, G. Lawrie, L. Grøndahl, European Polymer Journal 2016, 82, 1; b) F. Farhang, A. V. Nguyen, K. B. Sewell, Energy & Fuels 2014, 28, 7025.

[43] C. S. Goonasekera, K. S. Jack, J. J. Cooper-White, L. Grondahl, Journal of Materials Chemistry B 2013, 1, 5842.

[44] M. Soheilmoghaddam, H. Padmanabhan, J. J. Cooper-White, Biomaterials Science 2020, DOI: 10.1039/D0BM00946F.

