## Supplementary Material for "Driving Osteocytogenesis from Mesenchymal Stem Cells in Osteon-like Biomimetic Nanofibrous Scaffolds"

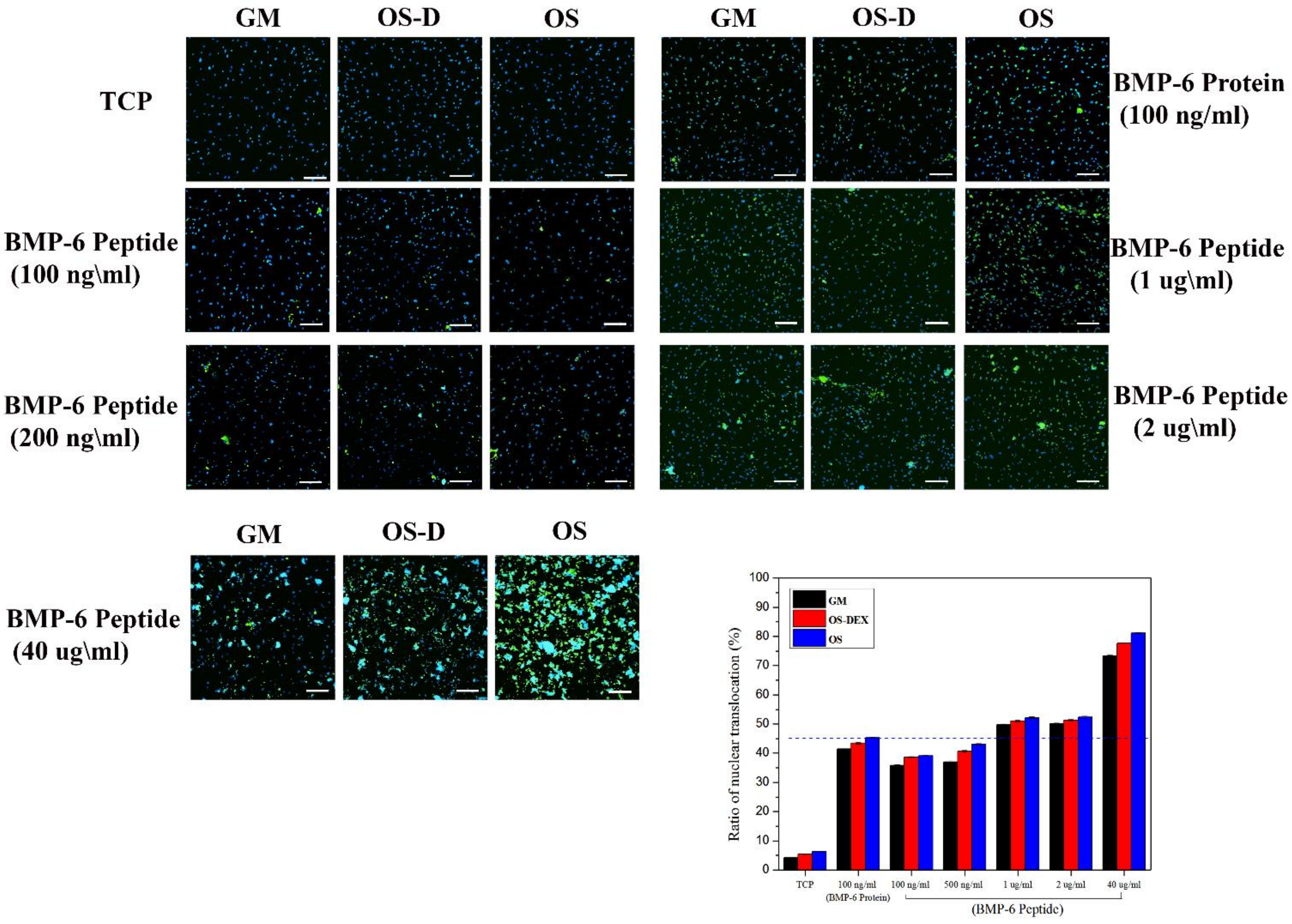


**Figure S 1.** Expression of P-Smad 1/5/8 induced by different dosages of BMP6 peptides (0, 100, 200, 1000, 2000 and 4000 ng/ml) after 24 hours. (a) P-Smad 1/5/8 and nuclei were detected by immunofluorescence using anti-phospho-Smad 1/5/8 antibody (green) and DAPI (blue), respectively. (b) The nuclear localisation of P-Smad 1/5/8 at different dosages of BMP6 peptides as well as 100 ng/ml of BMP6 protein was quantified. Scale bars, 200 μm; *P<0.05.

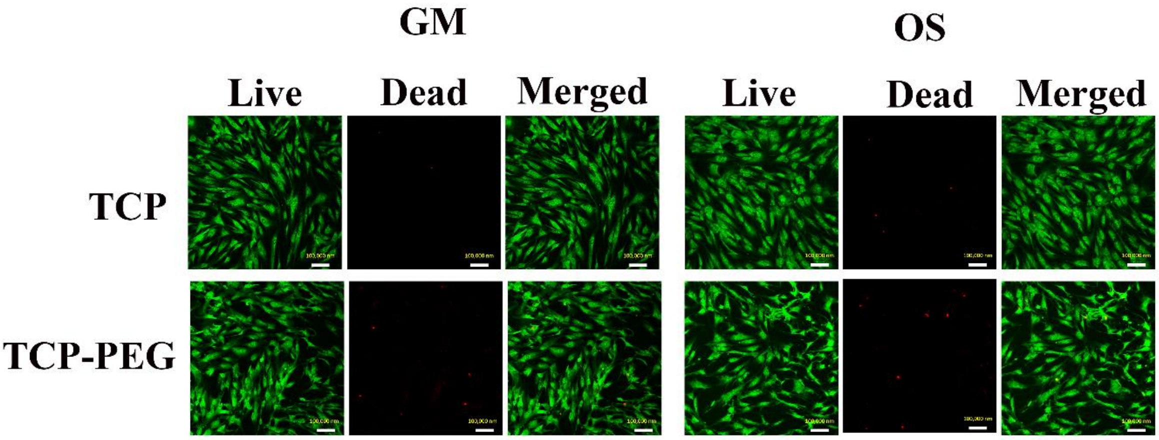


**Figure S 2.** Live/dead assay of hBMSCs covered with a layer of PEG-Nor/PEG-TH hydrogel. Green indicates live cells and red indicates dead cells. The scale bar indicates 100 μm.

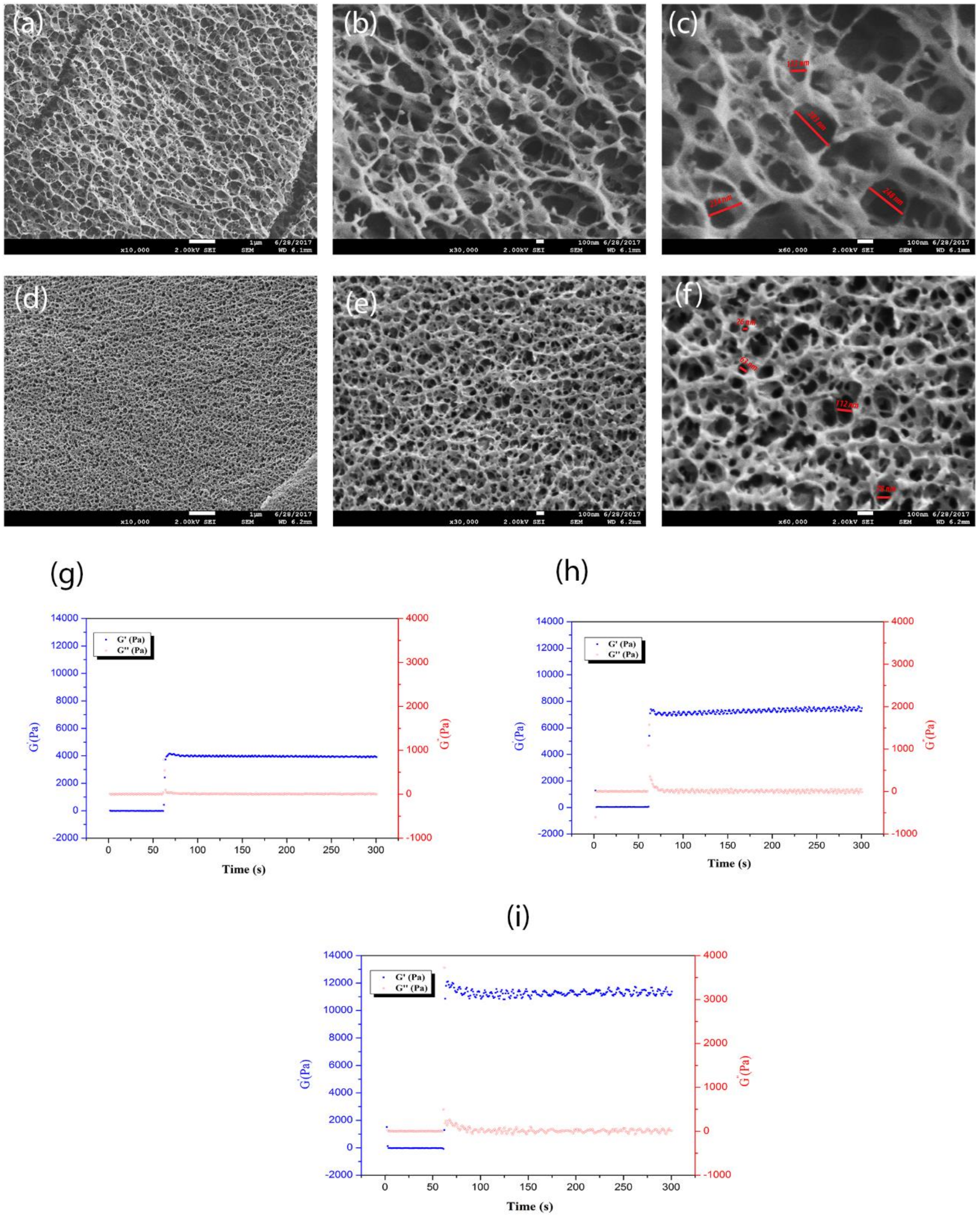


**Figure S 3**. Cryo-SEM pictures of (a, b and c) 4% (w/v) PEG-Nor/PEG-TH hydrogel, (d, e and f) 10% (w/v) PEG-Nor/PEG-TH hydrogel. Rheological Properties of (g) 4% (w/v) PEG-Nor/PEG-TH hydrogel, (h) 10% (w/v) PEG-Nor/PEG-TH hydrogel and (i) the BMP6 peptide conjugated 10% (w/v) PEG-Nor/PEG-TH hydrogel.

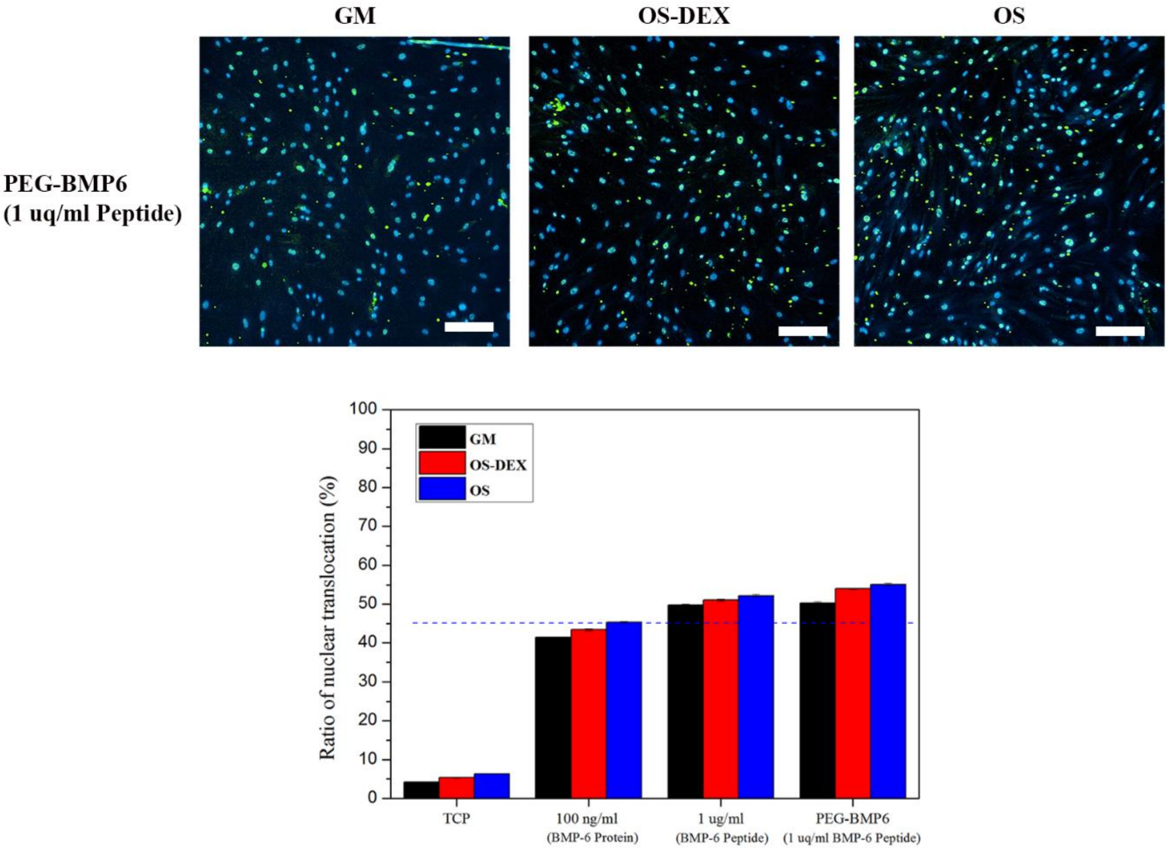


**Figure S 4.** Immunofluorescence staining of P-Smad 1/5/8 for hBMSCs cultured with BMP6 conjugated PEG hydrogel after 24 hours (1000 ng/ml of BMP6 peptide was used). The quantitative measurements of nuclei localised P-Smad 1/5/8 demonstrates that the BMP6 conjugated PEG enhanced P-Smad 1/5/8 nuclei localisation as compared to the cells that were exposed to BMP6 protein or same dosage of peptide in medium. P-Smad 1/5/8 is stained green and cell nuclei are stained blue. Scale bars, 200 μm; *$P<0.05$.

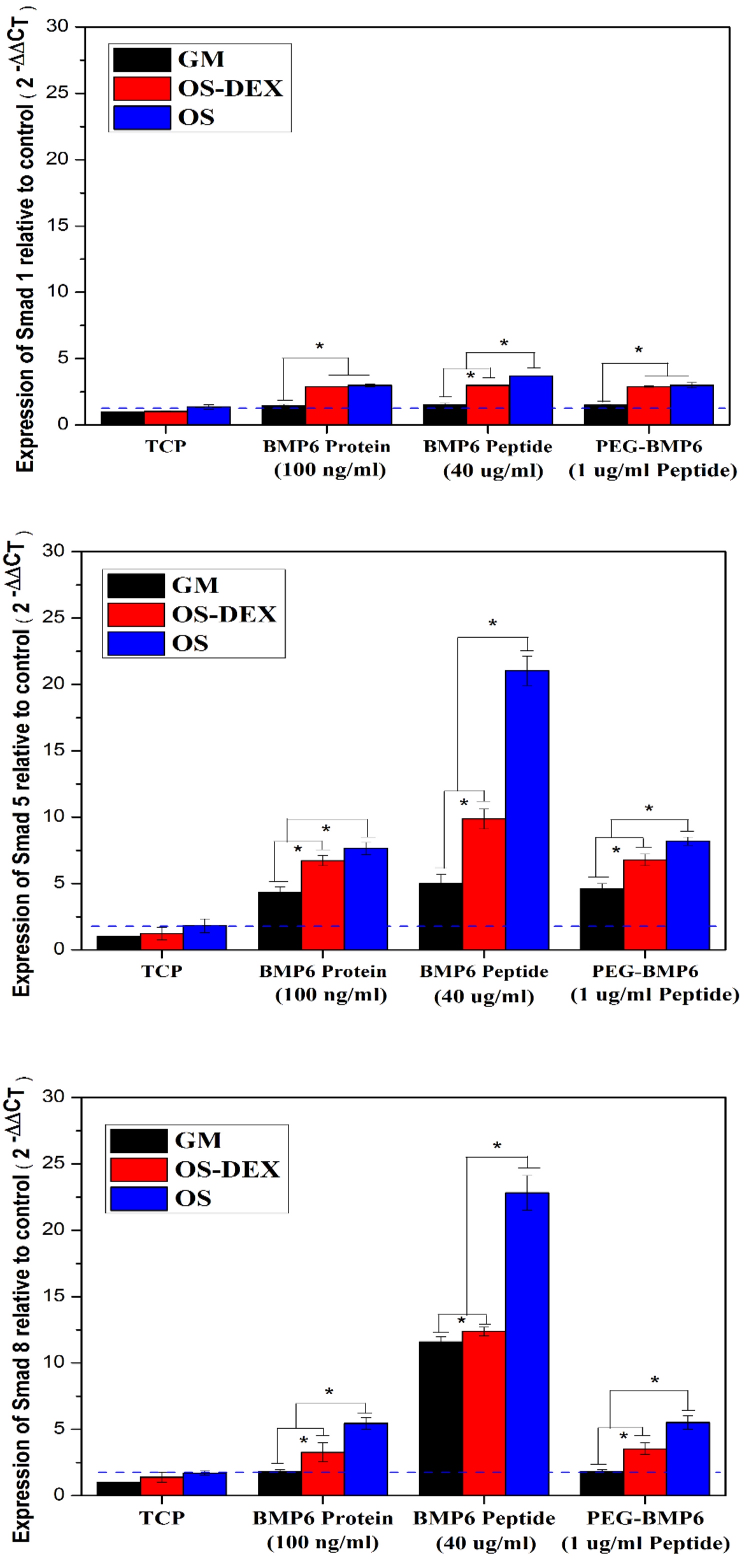


**Figure S 5.** Gene expression of Smad 1, Smad5 and Smad 8 for hBMSCs exposed to BMP6 protein and BMP6 peptide in medium or cultured with BMP6 conjugated PEG hydrogel after 24 hours. *$P<0.05$.

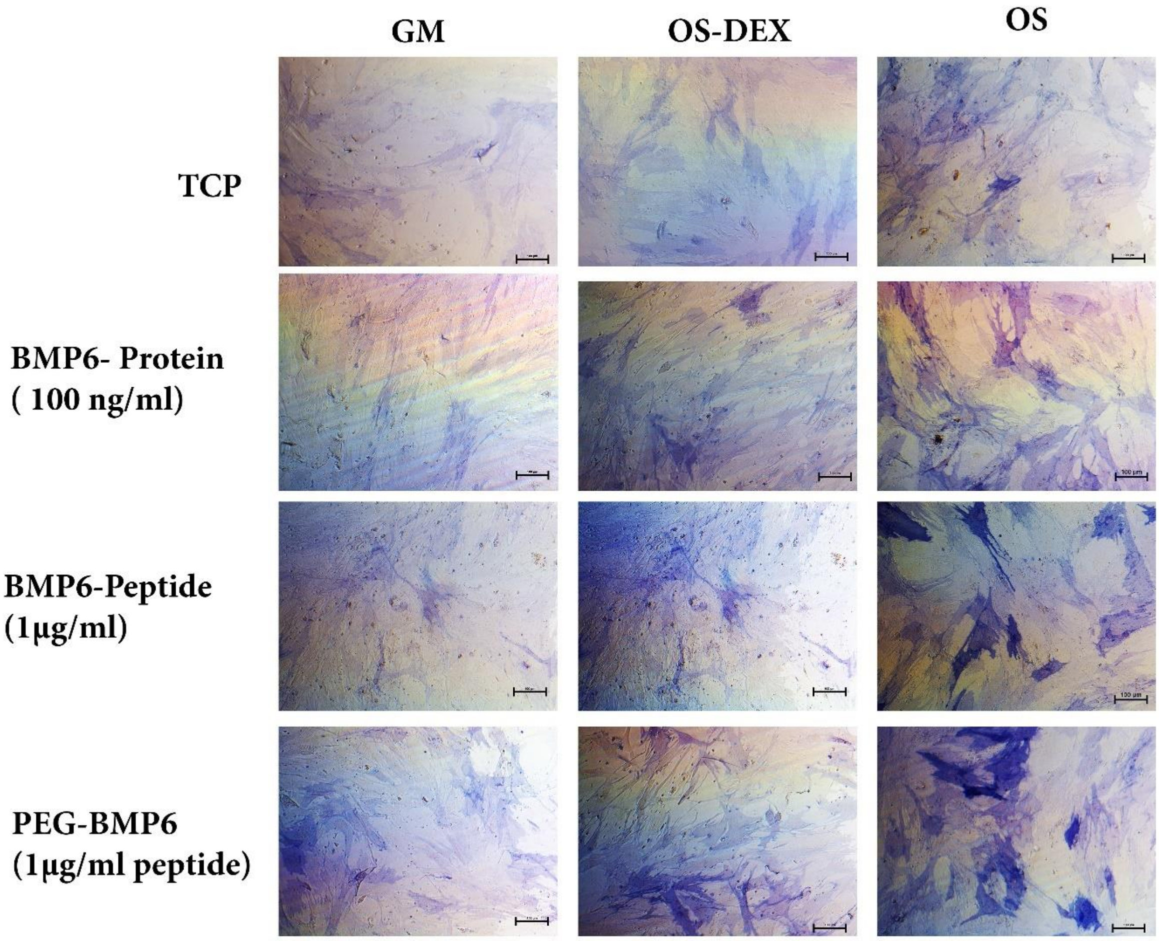


**Figure S 6.** ALP staining of the hBMSCs cultured for 7 days with 100 ng/ml BMP6 protein, 1000 ng/ml BMP6 peptide and BMP6 conjugated PEG-Nor/PEG-TH hydrogel with 1000 ng/ml BMP6 peptide dosage. Scale bar : 100 µm.

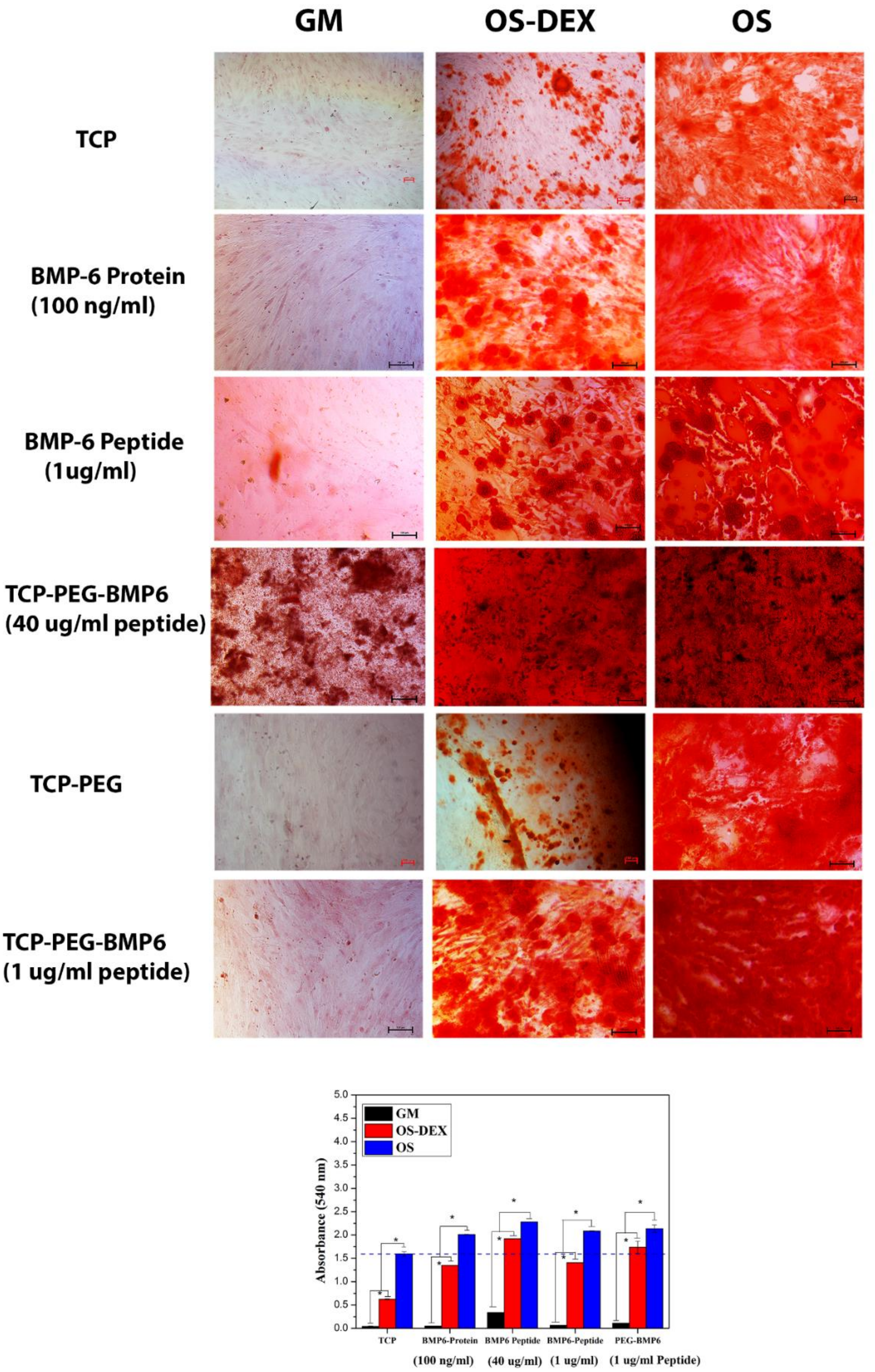


**Figure S 7.** Alizarin red staining results of hBMSCs cultured with BMP6 protein (100 ng/ml), different dosages of BMP6 peptides (1µg/ml and 40 µg/ml) and BMP6 conjugated PEG-Nor/PEG-TH hydrogels. The calcium deposition of the samples in different medium conditions was also quantified. Scale bars, 100 µm; *P<0.05.

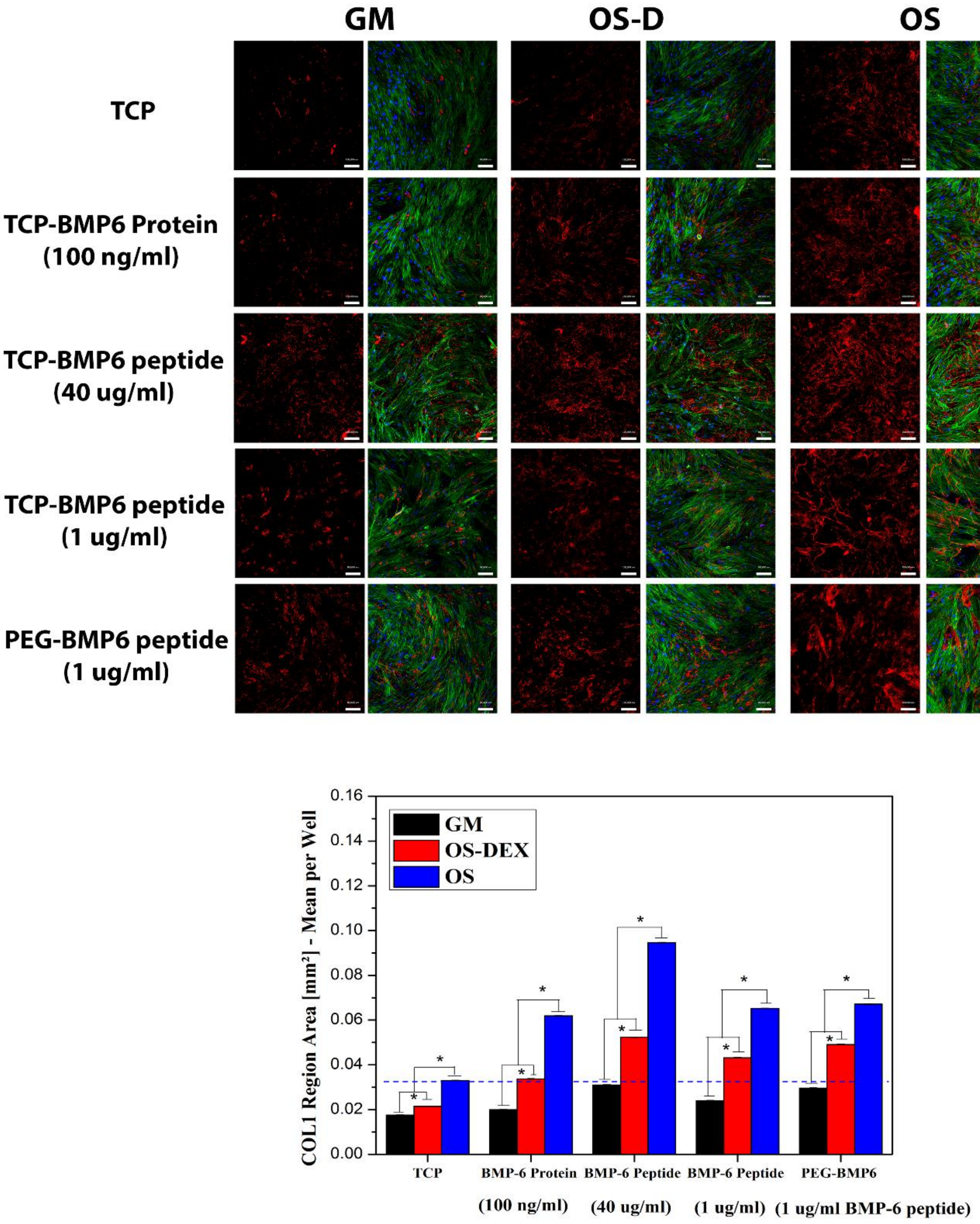


**Figure S 8.** Immunofluorescence of Col1 (in red) and nuclei (in blue) in hBMSCs after 7 days of exposure to BMP6 protein (100 ng/ml), different dosages of BMP6 peptides in medium (1 μg/ml and 40 μg/ml) and BMP6 peptide conjugated PEG hydrogels. Quantification of Col1 expression based on immunofluorescence staining was also conducted. Scale bars, 200 μm; *P<0.05.

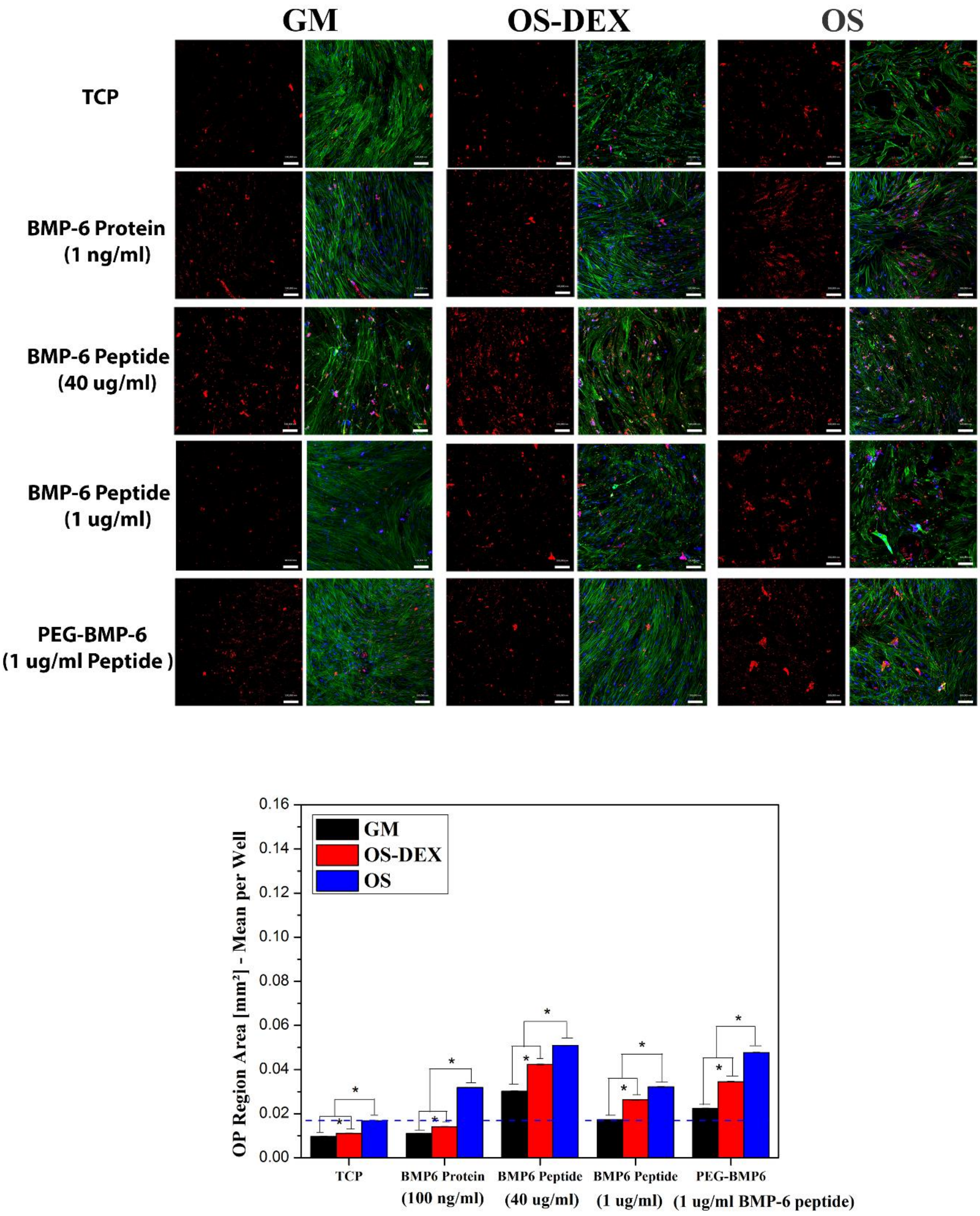


**Figure S 9.** Immunofluorescence of OP (in red) and nuclei (in blue) after hBMSCs were exposed to BMP6 protein, BMP6 peptide and BMP6 peptide conjugated into hydrogels for 21 days. Quantification of the area coverage by OP. Scale bars, 200 μm; *P<0.05.

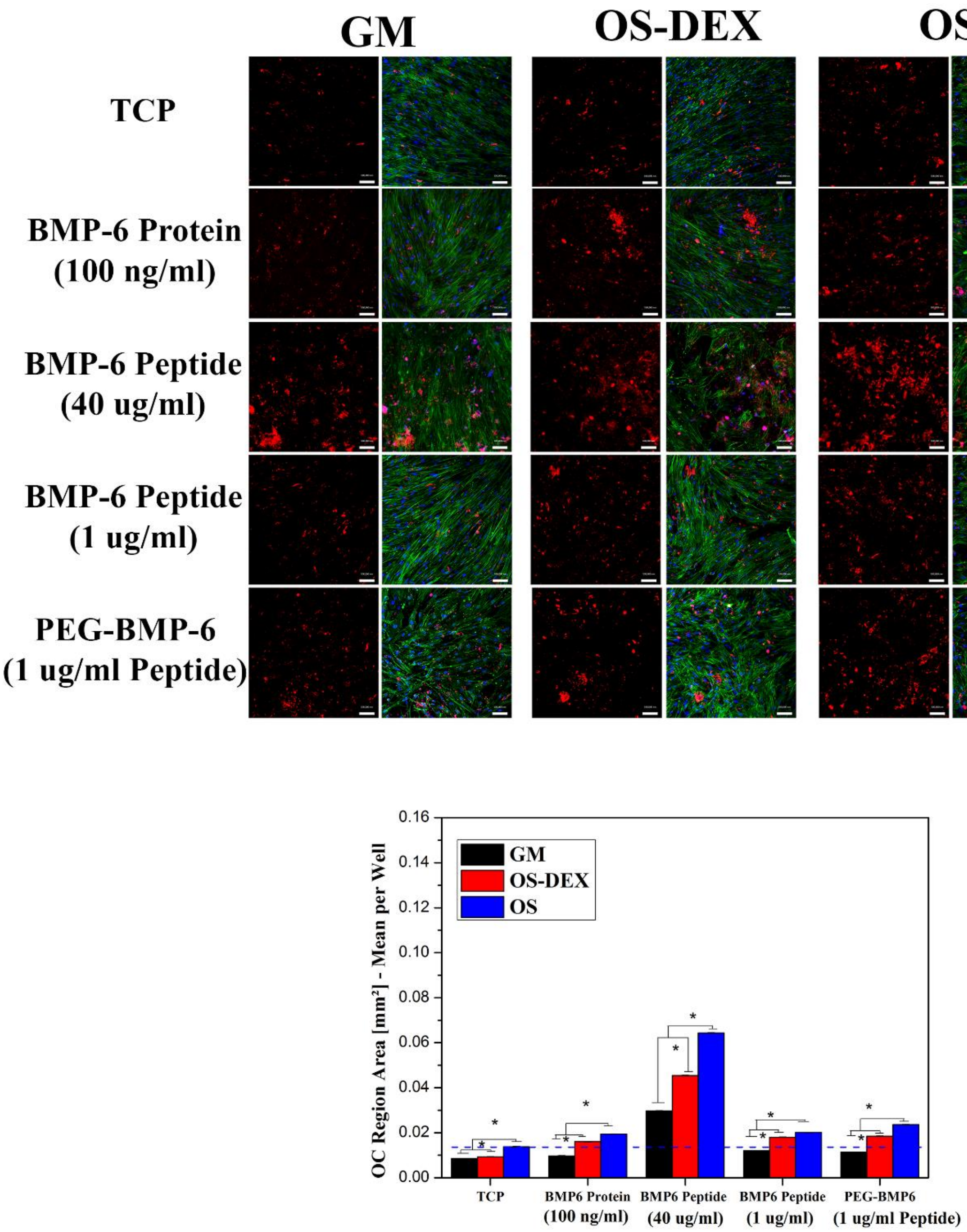


**Figure S 10.** Immunofluorescence staining of OC (in red) and nuclei (in blue) after hBMSCs were exposed to BMP6 protein, BMP6 peptide and BMP6 peptide conjugated into hydrogels for 21 days. Quantification of the area coverage by OC. Scale bars, 200 µm; *P<0.05.

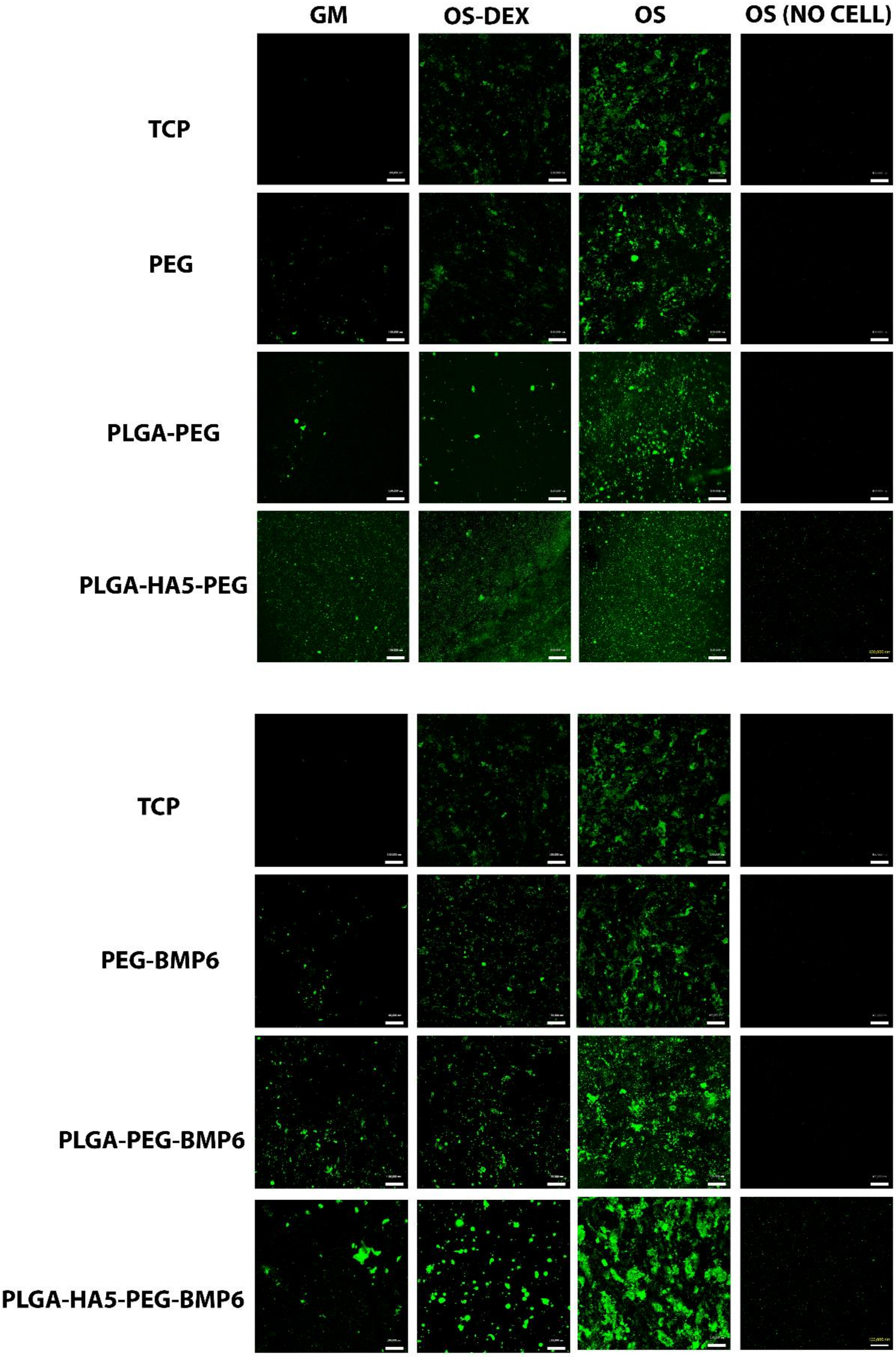


**Figure S 11.** Hydroxyapatite production on the bilayered scaffolds cultured in different medium conditions. Scale bar = 100 µm; $^{*}p < 0.05$

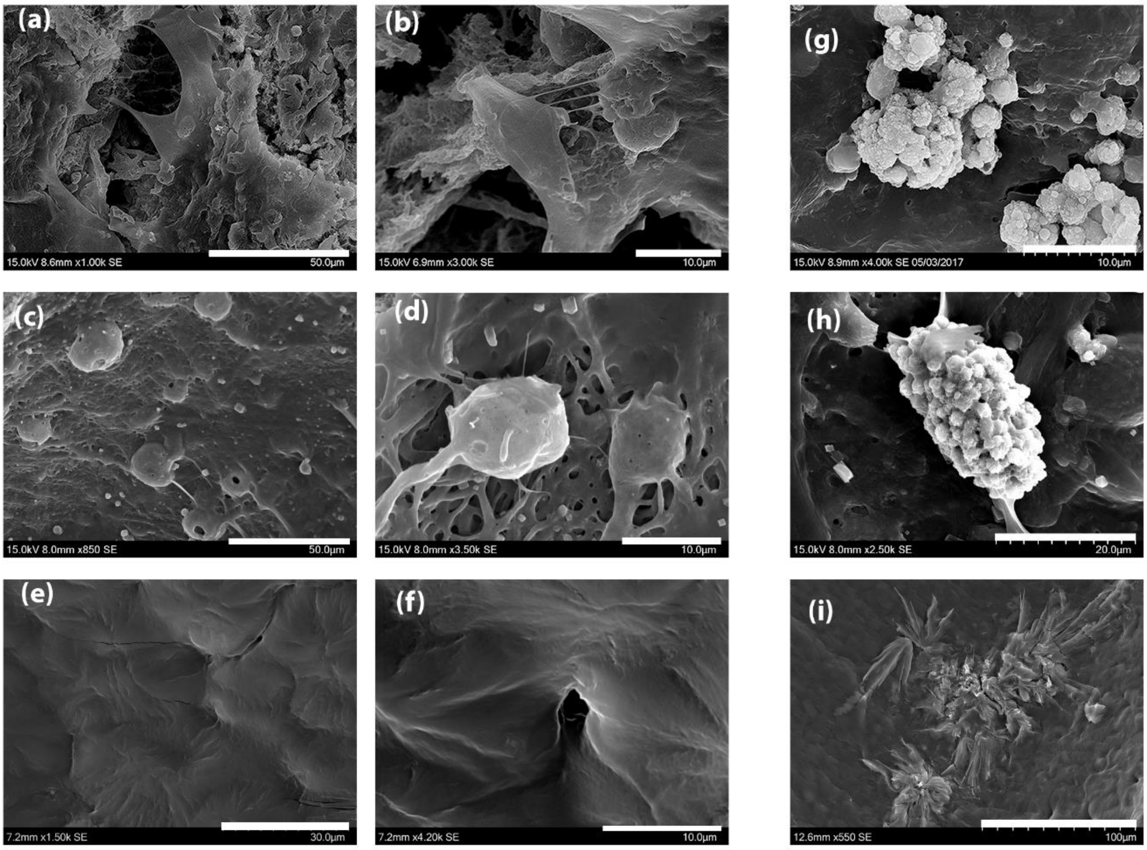


**Figure S 12.** SEM morphology of (a and b) mono-layered PLGA-HA5 electrospun mat, (c and d) PLGA-HA5-PEG bilayered scaffolds and (e and f) bilayered PLGA-HA5-PEG-BMP6 scaffold. The SEM pictures of new HA secreted on (g) mono-layered PLGA-HA5 electrospun mat, (h) bilayered PLGA-HA5-PEG and (i) bilayered PLGA-HA5-PEG-BMP6 scaffold.

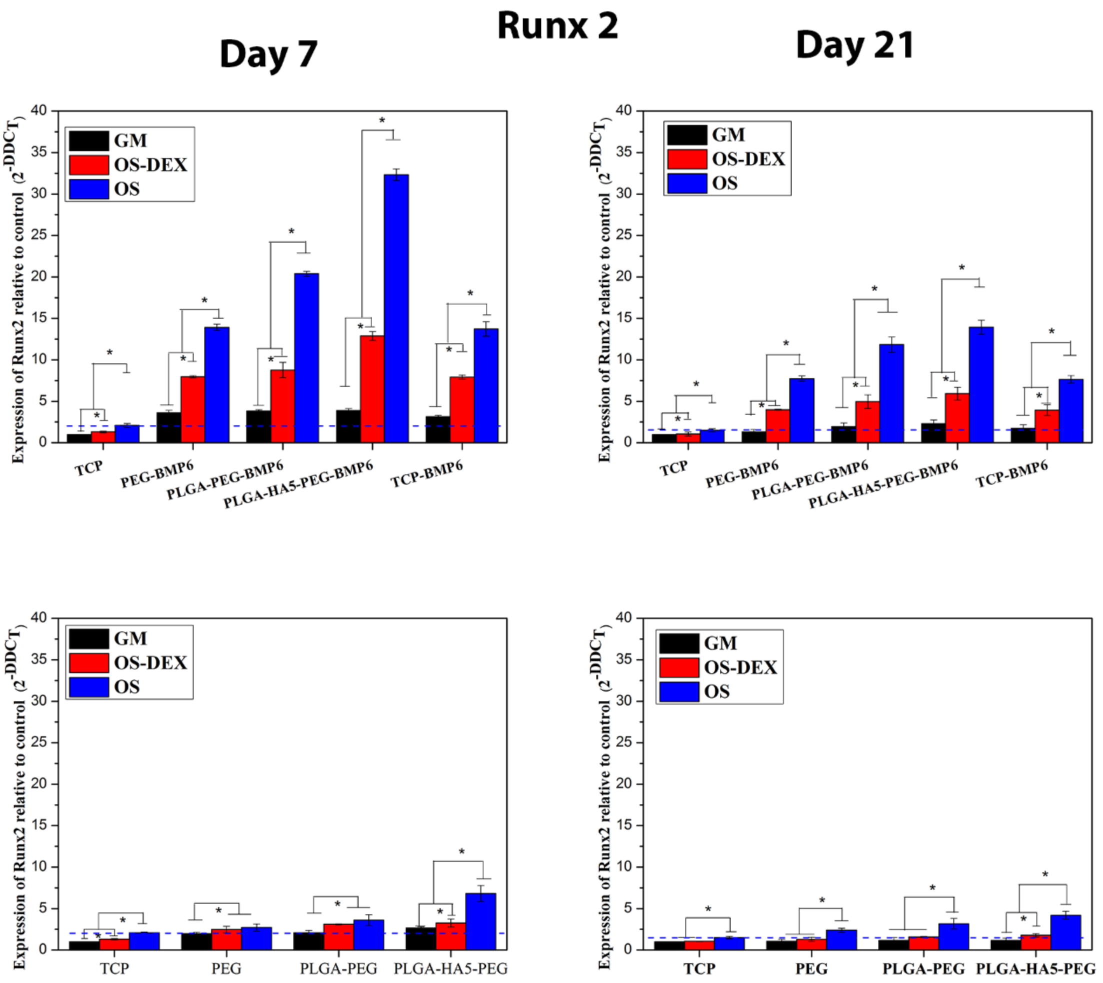


**Figure S 13.** Expression of the key transcript factors, Runx2, after 7 and 21 days of hBMSC cultivation on bilayered scaffolds in different medium conditions including GM, OS-DEX and OS. *P<0.05.

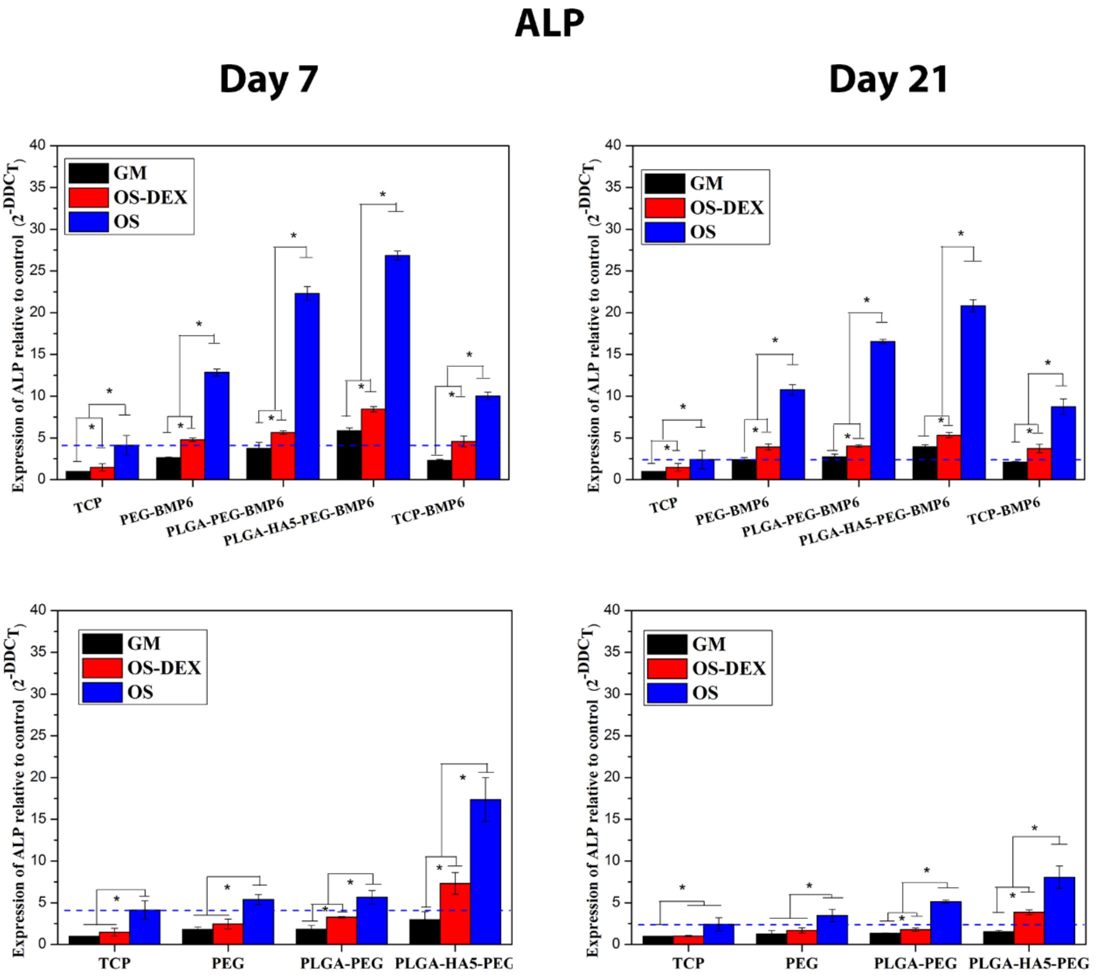


**Figure S 14.** Expression of early osteogenic marker ALP after 7 and 21 days of hBMSC cultivation on bilayered scaffolds in different medium conditions including GM, OS-DEX and OS. *$P<0.05$.

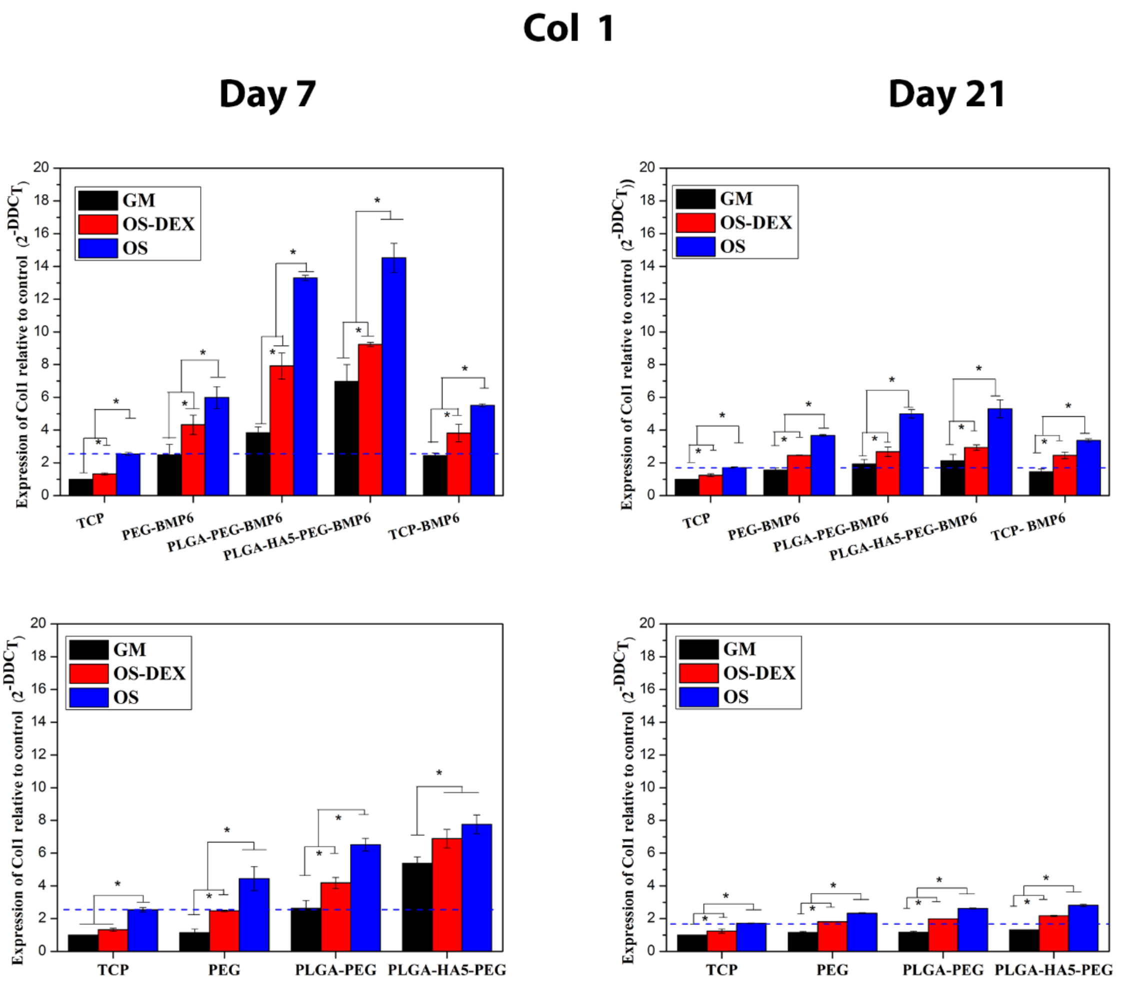


**Figure S 15.** Gene expression of Col1 for hBMSCs cultured on bilayered scaffolds in GM, OS-DEX and OS medium for 7 and 21 days. *P<0.05.

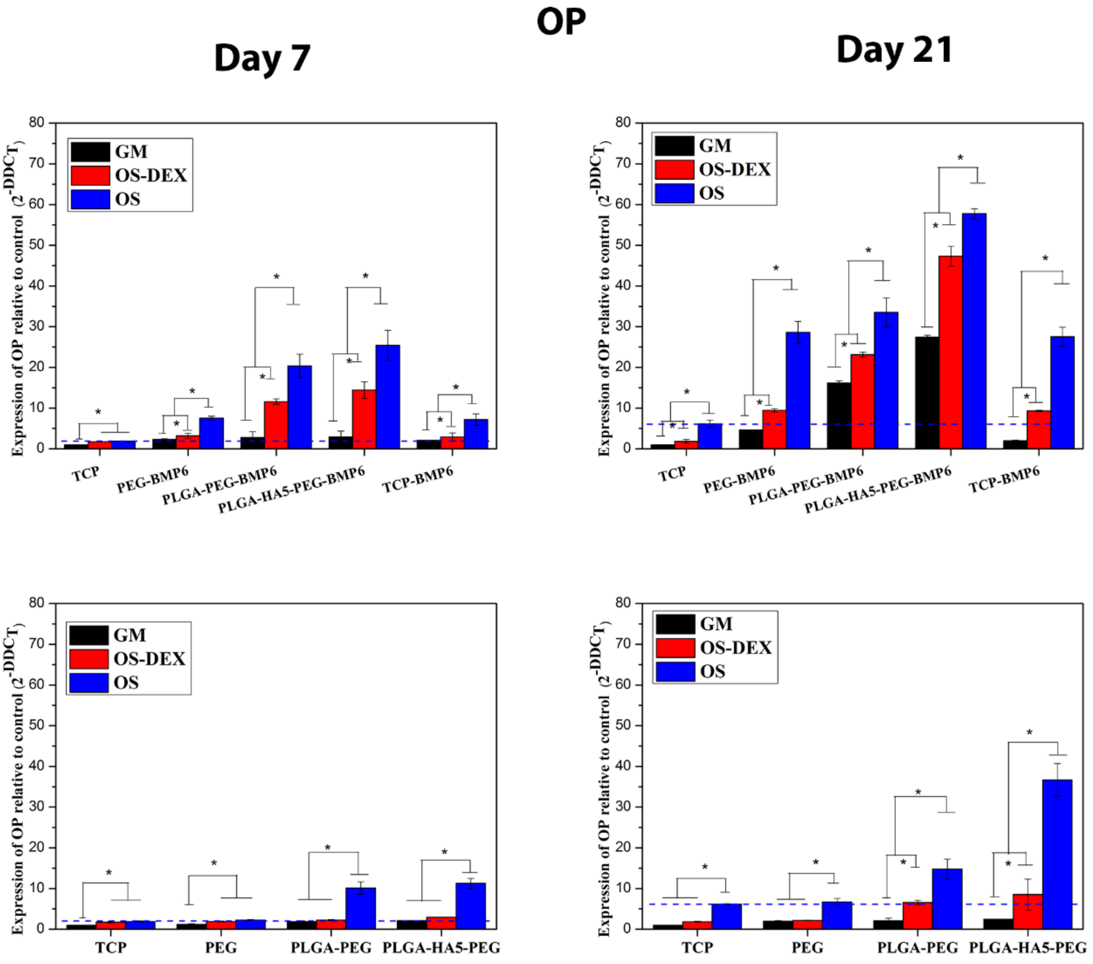


**Figure S 16.** The expression of late osteogenic gene OP was quantified by qPCR after 21 days of hBMSC cultivation on bilayered scaffolds in different medium conditions including GM, OS-DEX and OS. *P<0.05.

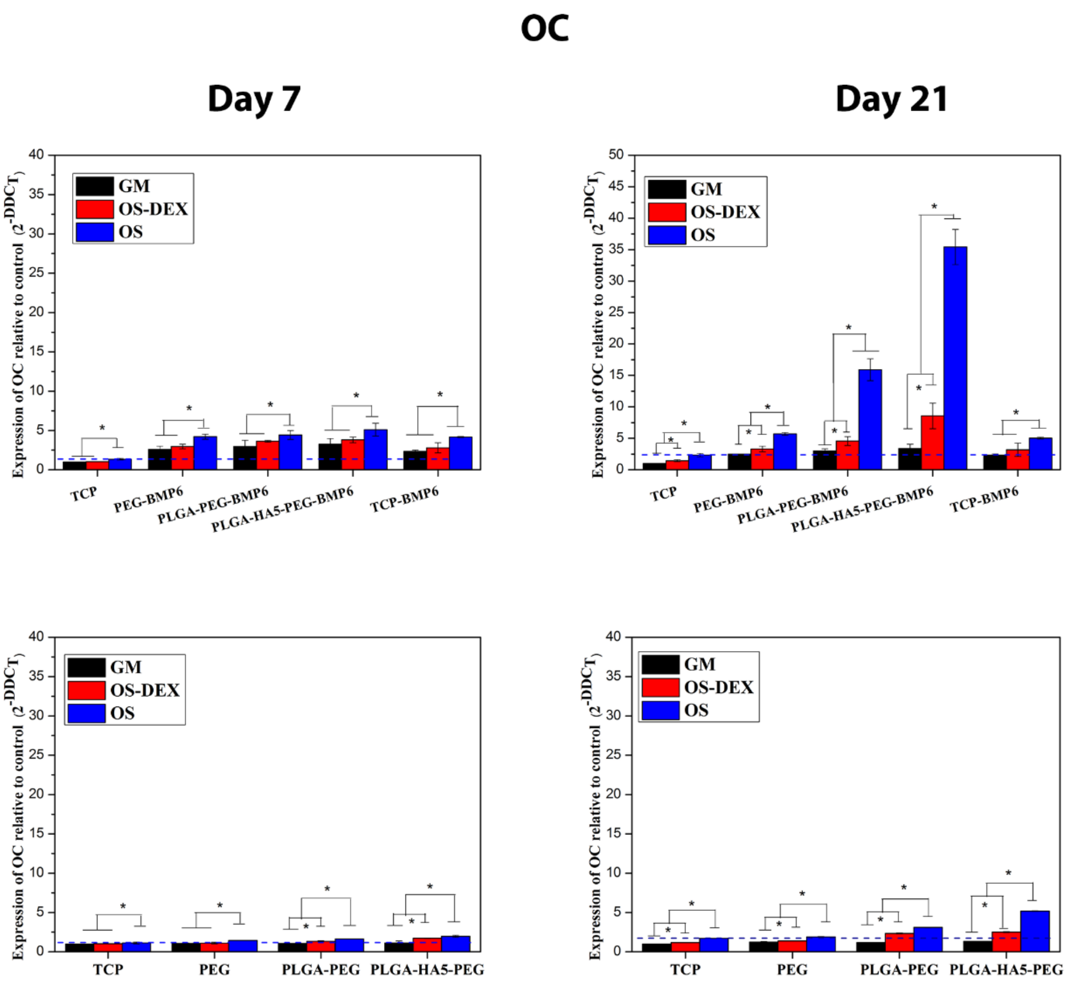


**Figure S 17.** The expression of mature bone marker OC was quantified by qPCR after 21 days of hBMSC cultivation on bilayered scaffolds in different medium conditions including GM, OS-DEX and OS. *P<0.05.

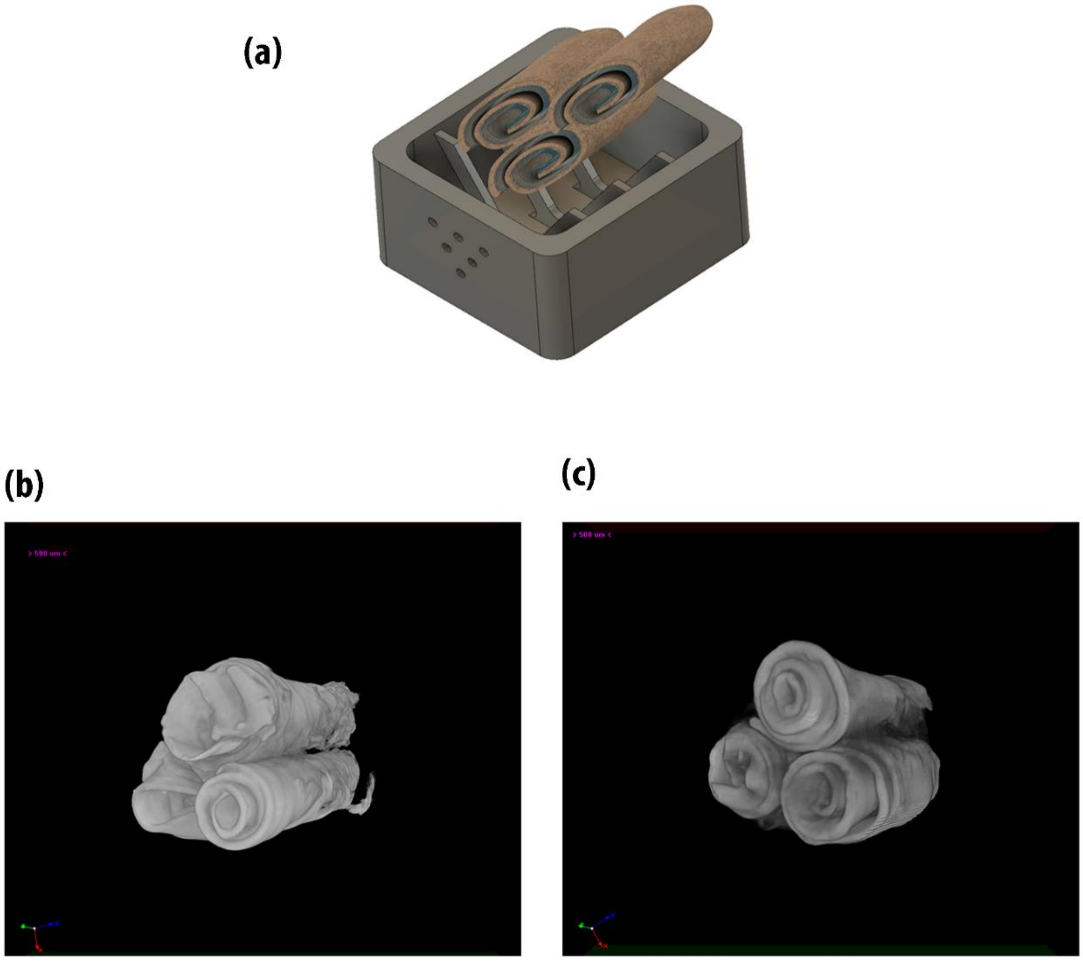


**Figure S 18.** (a) Incubation of osteon-like scaffolds inside a custom-designed bioreactor system where the osteogenic medium (GM and OS) flowed inside the bioreactor. (b) Micro CT image of osteon-like scaffolds in GM for 21 days. (c) Micro CT image of osteon-like scaffolds in OS for 21 days and. Scale bar : 500μm

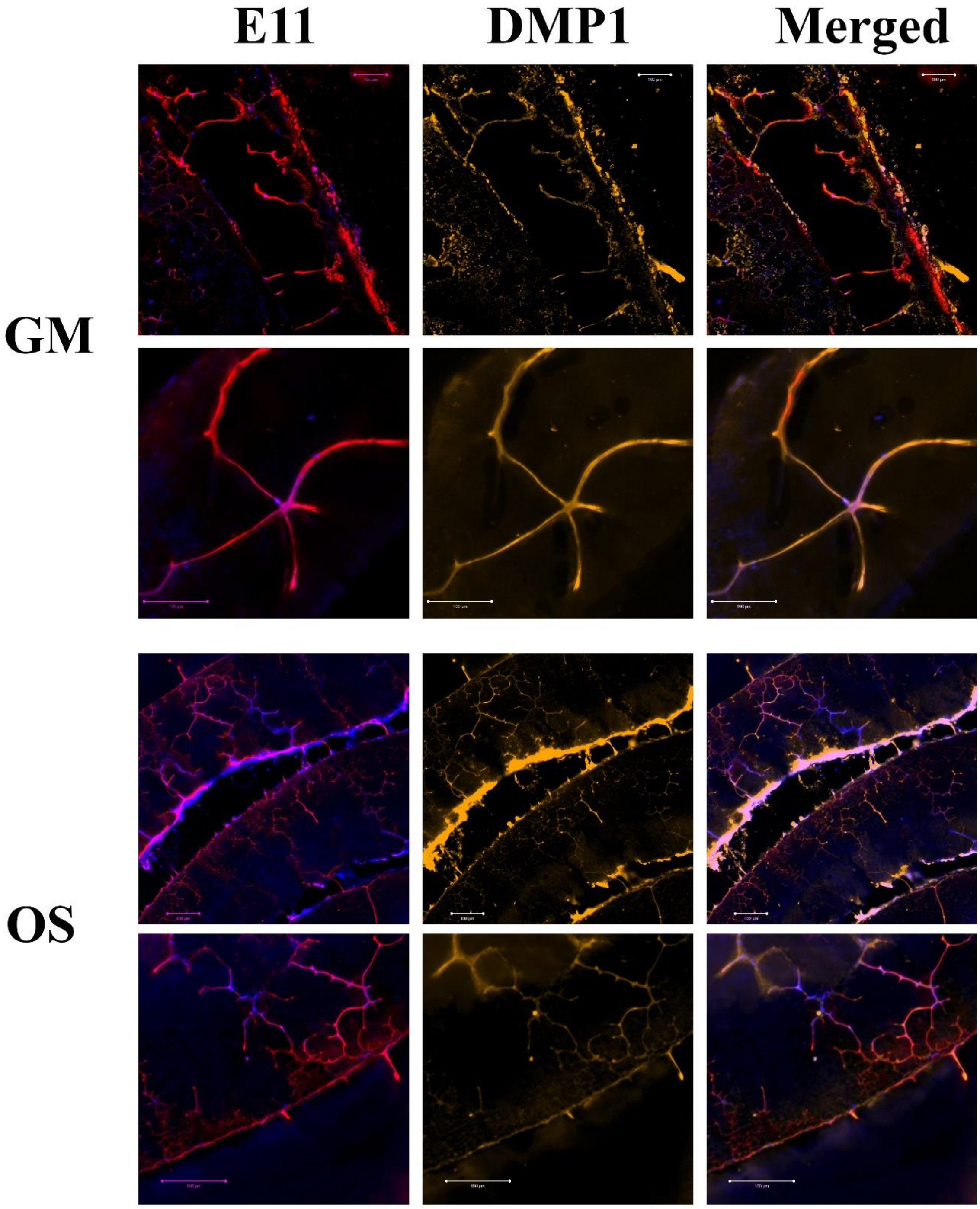


**Figure S 19.** Immunofluorescence staining of cross-sectional view of osteon-like samples cultured in bioreactor for 21 days. The samples were stained positive for E11(red) and DMP1 ( yellow) forming a multiple branching dendrites morphology that migrated into the neibouring layers. The merged pictures represents E11, DMP1 and nuclei (blue).

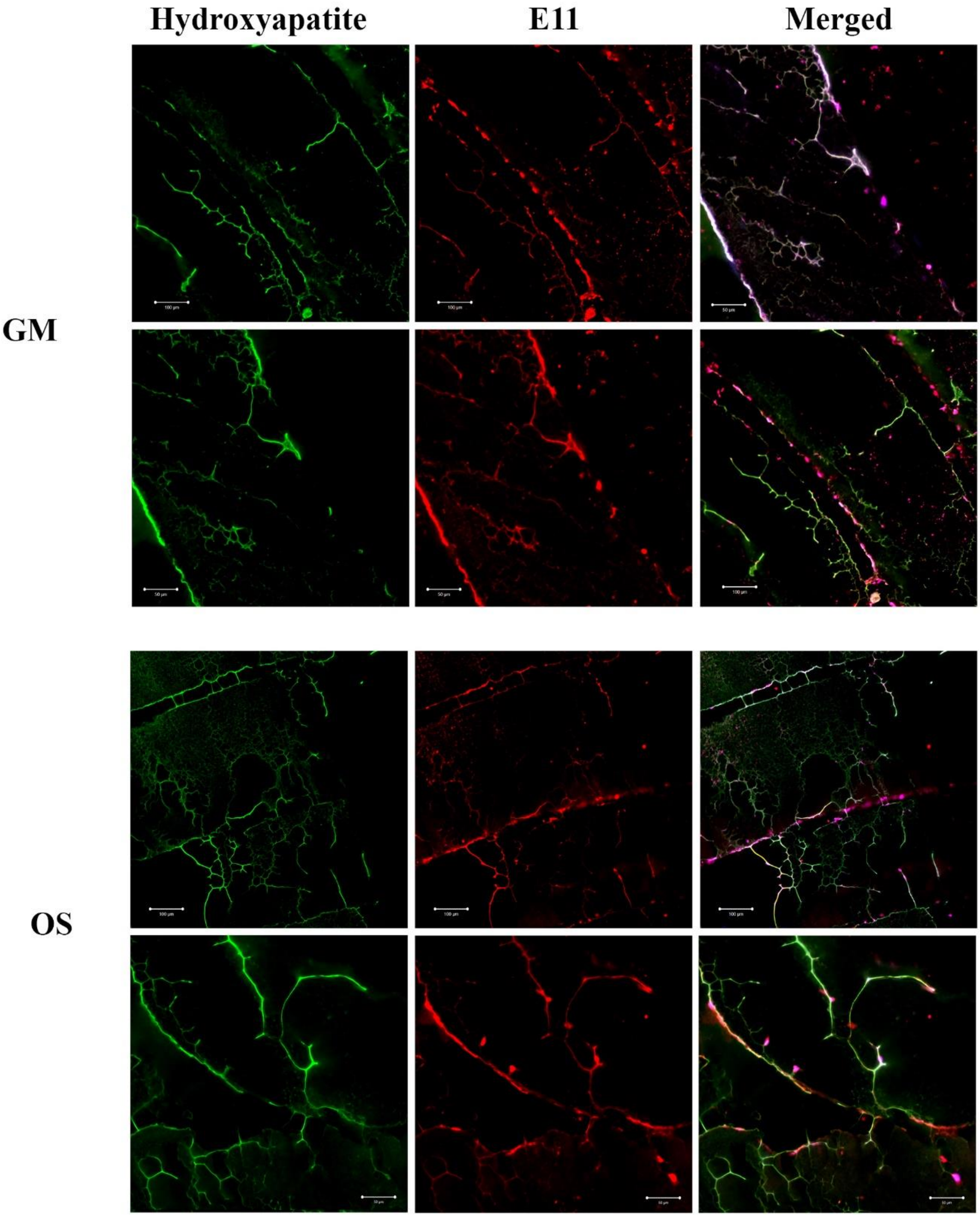


**Figure S 20.** Immunofluorescence staining of cross-sectional view of osteon-like samples cultured in bioreactor for 21 days. The samples were stained positive for hydroxyapatite (green) E11(red). The merged pictures represents hydroxyapatite, E11 and nuclei (blue)

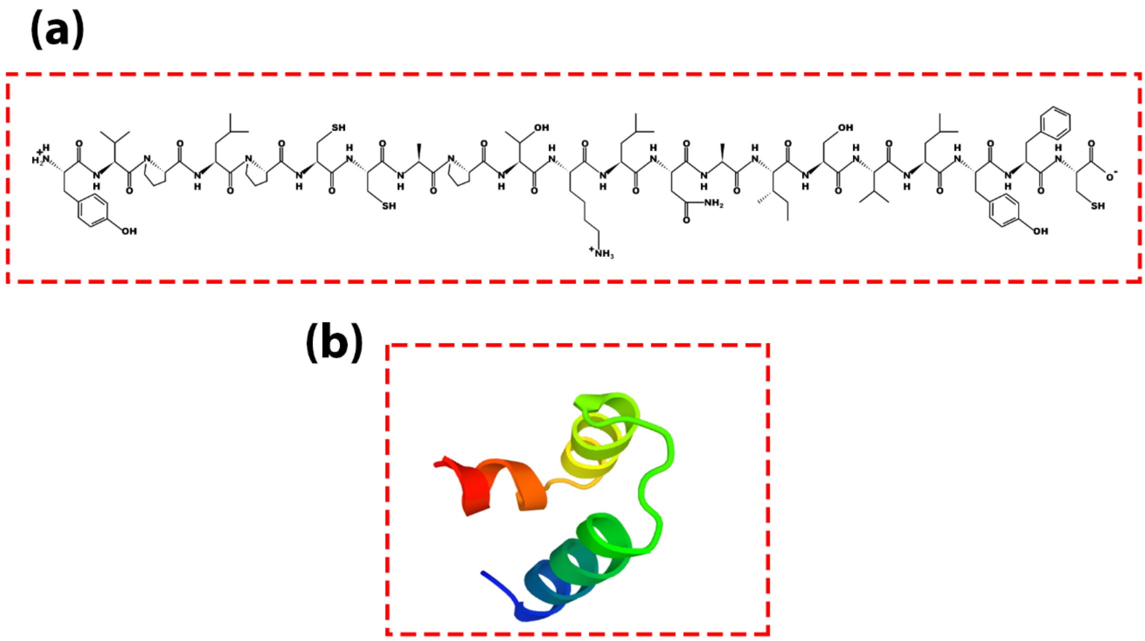

**Figure S 21.** (a) Chemical structure of the synthesised cysteine -terminated BMP6 peptide and molecular structure of the BMP6-derived peptide.

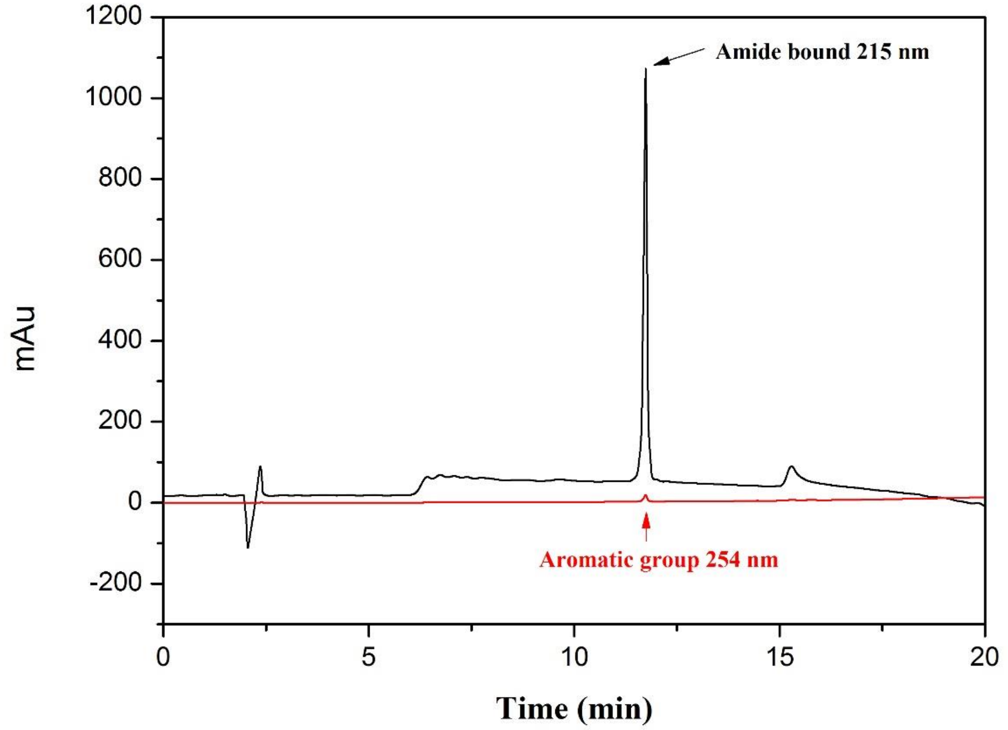


**Figure S 22.** HPLC data obtained for the synthesised BMP6 peptide.

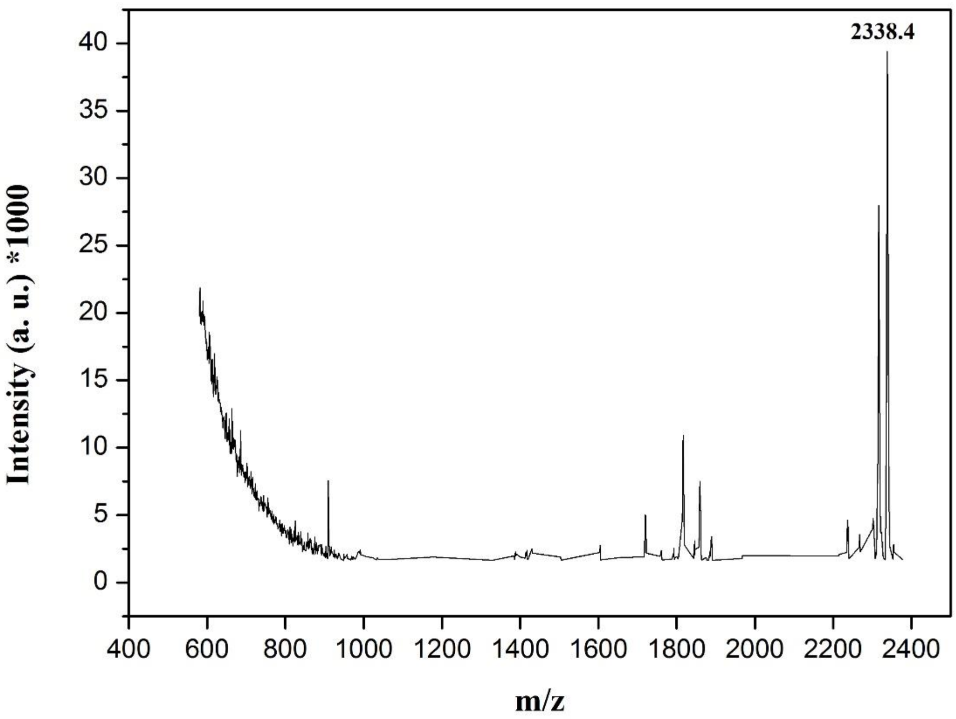


**Figure S 23.** Matrix-assisted laser desorption/ionisation (MALDI) data obtained for the synthesised BMP6 peptide.
